# Spatial Effects of Infiltrating T cells on Neighbouring Cancer Cells and Prognosis in Stage III CRC patients

**DOI:** 10.1101/2024.01.30.577720

**Authors:** Mohammadreza Azimi, Sanghee Cho, Emir Bozkurt, Elizabeth McDonough, Batuhan Kisakol, Anna Matveeva, Manuela Salvucci, Heiko Dussmann, Simon McDade, Canan Firat, Nil Urganci, Jinru Shia, Daniel B. Longley, Fiona Ginty, Jochen H. M. Prehn

## Abstract

Colorectal cancer (CRC) is one of the most frequently occurring cancers, but prognostic biomarkers identifying patients at risk of recurrence are still lacking. In this study, we aimed to investigate in more detail the spatial relationship between intratumoural T cells, cancer cells, and cancer cell hallmarks, as prognostic biomarkers in stage III colorectal cancer patients. We conducted multiplexed imaging of 56 protein markers at single cell resolution on resected fixed tissue from stage III CRC patients who received adjuvant 5-fluorouracil-based chemotherapy. Images underwent segmentation for tumour, stroma and immune cells, and cancer cell ‘state’ protein marker expression was quantified at a cellular level. We developed a Python package for estimation of spatial proximity, nearest neighbour analysis focusing on cancer cell – T cell interactions at single-cell level. In our discovery cohort (MSK), we processed 462 core samples (total number of cells: 1,669,228) from 221 adjuvant 5FU-treated stage III patients. The validation cohort (HV) consisted of 272 samples (total number of cells: 853,398) from 98 stage III CRC patients. While there were trends for an association between percentage of cytotoxic T cells (across the whole cancer core), it did not reach significance (Discovery cohort: p = 0.07, Validation cohort: p = 0.19). We next utilized our region-based nearest neighbourhood approach to determine the spatial relationships between cytotoxic T cells, helper T cells and cancer cell clusters. In the both cohorts, we found that lower distance between cytotoxic T cells, T helper cells and cancer cells was significantly associated with increased disease-free survival. An unsupervised trained model that clustered patients based on the median distance between immune cells and cancer cells, as well as protein expression profiles, successfully classified patients into low-risk and high-risk groups (Discovery cohort: p = 0.01, Validation cohort: p = 0.003).

## Introduction

Colorectal cancer (CRC) shows heterogeneous clinical response, and prognostic biomarkers for identifying people at elevated risk of disease recurrence are still underdeveloped [1]. In cases where surgery (and/or neoadjuvant chemoradiation therapy in the rectal cancer setting) may not completely eliminate cancer cells, chemotherapy can be used as adjuvant therapy to reduce the risk of cancer recurrence [2]. The primary goal of adjuvant chemotherapy is to kill or inhibit the growth of micrometastatic cancer cells. In stage III CRC, tumour cells starts to spread into surrounding tissues and lymph nodes. Adjuvant, 5-Fluorouracil-based treatment regimens are standard-of-care (SOC) for the management of stage III CRC with lymph node involvement [3].

The tumour microenvironment and specifically immune responses are an emerging component for cancer recurrence prediction. The tumour microenvironment is a complex milieu where interactions between cancer cells and the immune system play a pivotal role in shaping disease progression and therapy responses [4]. CD8^+^ or cytotoxic T cells are known for their antitumour immune response, releasing perforin and granzymes to create pores in the tumour cell plasma membrane and initiate cell death [5, 6]. However, not all CD8^+^ T cells are equally effective in cell killing, due to exhaustion, high expression of checkpoint proteins in cancer cells, and both cell number and distribution within the tumour can affect patient outcomes. In a variety of tumour types, including, CRC, breast cancer, and non-small cell lung cancer, high numbers of infiltrating cytotoxic T cells are correlated with good prognosis [5–12]. The association between increased CD8^+^ T cell density and a favourable outcome, however, does not apply to all cancer types. For instance, in the case of bladder cancer, high numbers of infiltrating CD8^+^ and TIGIT^+^ (marker of exhaustion) cells are associated with poor prognosis [13] and in renal cell cancer (RCC), higher densities of CD8^+^ with checkpoint protein expression (PD-1, LAG-3, PD-L1, and PD-L2) correlated with poor prognosis [14]. Further, the proximity of functional CD8^+^ T-cells to cancer cells is essential to elicit their cytotoxic effects [15–17].

CD8^+^ T cells can recognize and target cancer cells for destruction through various mechanisms, including inducing apoptosis [17]. Expression of HLA-1 antigens and apoptosis-regulating proteins in cancer cells may influence the interaction efficacy between CD8^+^ T cells and cancer cells by regulating immune recognition and cell death susceptibility, respectively. It is not yet fully elucidated whether cancer cells that are closer to CD8^+^ T cells exhibit altered expression of apoptosis proteins, HLA-1 class antigens, or other cancer cell hallmarks such as cell proliferation and cell metabolism markers, which may potentially impact on cancer cell recognition by CD8^+^ T cells and subsequently on patient outcomes.

We here employed a multiplexed immunofluorescence imaging (Cell DIVE) technique [18] that allows for the simultaneous measurement of multiple biomarkers and colocalization analysis on a single fixed 5µm tissue section. These techniques provide spatial context, allowing us to visualize the distribution of CD8^+^ T cells in relation to tumour cells, stromal cells, and other components of the tumour microenvironment, and reveal the spatial and phenotypic heterogeneity of CD8^+^ T cells and cancer cells within the tumour.

## Methods and Materials

Figure.1 depicts the workflow for this study. We have applied the following approaches to evaluate the prognostic significance of the median distance between intratumoural T cells and cancer cells across discovery and validation cohorts.

**Figure. 1.**
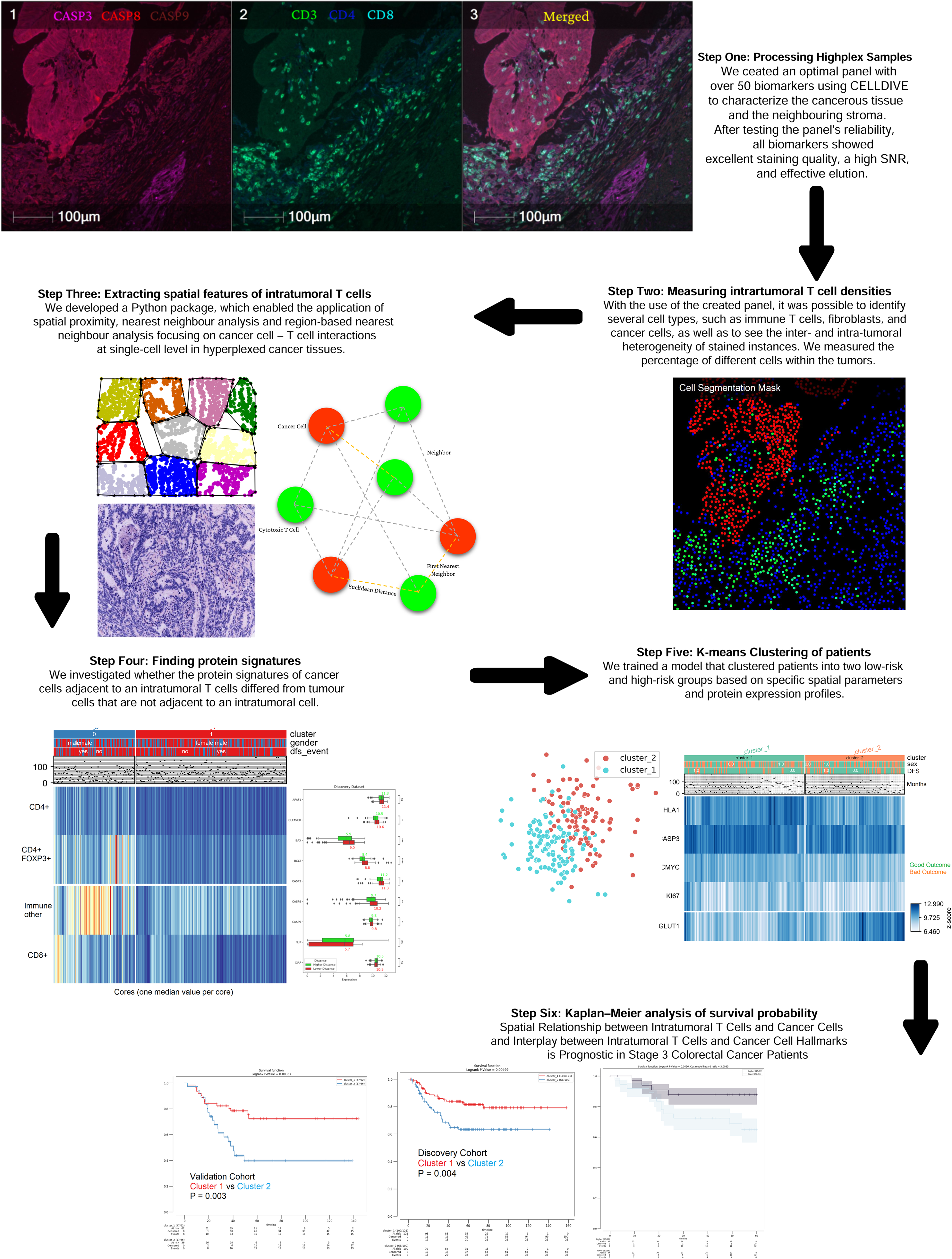
Workflow: We created an optimal panel with over 50 biomarkers using CELLDIVE to characterize the cancerous tissue and the neighbouring stroma. After testing the panel’s reliability, all biomarkers showed excellent staining quality, a high SNR, and effective elution. With the use of the created panel, it was possible to identify several cell types, such as immune T cells, fibroblasts, and cancer cells, as well as to see the inter- and intra-tumoural heterogeneity of stained instances. We obtained nearest neighbourhood analysis data and tested the prognosis of the cancer hallmarks. 1): Mutliplexed imaging of Samples, 2): Measuring intratumoural T cell densities; 3): Extracting spatial features of intratumoural T cells; 4): Finding protein signatures associated with proximity to immune cells; 5): K-means Clustering of patients Six: Kaplan–Meier analysis of survival probability

The total number of patients in the discovery dataset was 221 (n=121 females and 100 males). For construction of the tissue microarray (TMA), three cores (1 mm in diameter) from the tumour center of formalin fixed paraffin embedded (FFPE) samples were selected by a pathologist (Dr. Jinru Shia, Memorial Sloan Kettering Cancer Center) for each patient. A total of 1627 cores were selected and 8 tissue blocks were constructed. The discovery dataset (MSK) consisted of 462 multiplex immunofluorescence images of stage III CRC patients and the total number of segmented cells after QC was 1,669,228 (QC workflow is presented in **Supplemental Figure 1**).

The validation cohort was collected from the Huntsville Clearview Cancer Center (HV) and consisted of 272 FFPE tumour cores (three cores per patient) from 98 stage III CRC patients with adjuvant therapy. The clinical and pathological variables are presented in **Supplemental Table 1**. The training and validation cohorts were comparable and no statistical differences in clinical variables were found.

The TMA/clinical data QC exclusion criteria included: i) poor tissue quality or no tumour cells in tissue/adjacent normal samples; ii), heavily artefacted tissue, extensive tumour necrosis, extensive presence of normal adjacent tissue; iii) loss of follow-up or recurrence and/or death within less than two months from surgical resection; iv) stage II or IV disease; v) rectal or mucinous tumours. After applying exclusion criteria from the original patient cohort, we selected patients who were only treated with 5FU-based adjuvant chemotherapy (n discovery= 221 stage III patients and n validation =98).

This study was performed in accordance with ethical guidelines for clinical research with the approval of the ethics committee for each institution (Memorial Sloan Kettering Cancer Center and Huntsville Clearview Cancer Center). Between January 1997 and December 2008, all of the study’s participants had surgery at Memorial Sloan Kettering Cancer Center or Huntsville Clearview Cancer Center for stage III CRC that was confirmed by histology.

Multiplexed immunofluorescence imaging (MxIF) (Cell DIVE, Leica Microsystems, Issaquah) was used to analyze TMAs as previously described by Gerdes et al [19]. Briefly, TMAs were de-paraffinized and rehydrated, underwent a two-step antigen retrieval and were then stained for 1 hour at room temperature using a Leica Bond autostainer. Antibodies against DAPI, pan-cytokeratin (CK-26), ribosomal S6, and NaKATPase were used to separate epithelial and stromal cells. For the objective of this study, we focused on expression of immunological markers: CD3, CD4, CD8, CD20, forkhead box P3 (FOXP3), and cancer and cancer-related hallmarks: HLA class 1 (HLA-1), C-myc, GLUT-1, Caspase-3, caspase-8, caspase-9, BAK, BAX, FLIP, FADD, APAF1, GRP78, BCL2, XIAP, HK2, LDHA, PKM2, Ki67, and TIGAR. All antibodies were characterized per the previously described protocol [19]. After down-selection, each antibody was conjugated with either Cy3 or Cy5 bis-NHS-ester dyes using standard protocols as previously described by Gerdes et al [19]. All samples underwent DAPI imaging in every round, and background (inherent tissue autofluorescence prior to staining) imaging for the first five rounds and every three rounds thereafter.

Using Cell DIVE automated image pre-processing software, all images were registered to baseline using DAPI and underwent background autofluorescence subtraction, illumination and distortion correction. DAPI and Cy3 autofluorescence images were used to generate a pseudo-colored image, which visually resembles a hematoxylin and eosin (H&E) stained image, which we refer to as a virtual H&E (vH&E). This visualization format helps tissue QC review and facilitated review of tumour morphology and lymphocytes. All cells in the epithelial and stromal compartments were segmented using DAPI and CK-26, while S6, and NaKATPase were used for subcellular analysis of epithelial cells. Each segmented cell was assigned an individual ID and spatial coordinate. Post segmentation, several quality control (QC) steps were conducted (described in detail in Berens et al [20]), including visual review and manual scoring of tissue quality and segmentation for every image. Each image was reviewed for completeness and accuracy of segmentation masks in each subcellular compartment and tumour and stroma separation. Average biomarker intensity was calculated for each cell and the following additional cell filtering criteria were applied: 1) epithelial cells were required to have either 1-2 nuclei; 2) each sub-cellular compartment (nucleus, membrane, cytoplasm) area had to have > 10 pixels and < 1500 pixels; 3) cells had to have excellent alignment with the first round of staining (round 0) ; 4) cells were at >25 pixels distance from the image margins; 5) cell area for nuclear segmentation mask was >100 or <3000 pixels, 6) duplicates.

For immune cell classification a machine learning approach with an automatically generated annotated training set was applied [21]. Briefly, the auto-fluorescence-removed images were segmented at cellular level to identify cells that were potentially positive for each immune marker (CD3^+^, CD4^+^, CD8^+^, CD20^+^, FOXP3^+^ and PD1^+^) via cellular intensity and morphological criteria. These candidate annotations were then correlated with segmented nuclei and segmented cells with no nuclei were discarded. The remaining positive cells (or “auto-annotated cells”) were the automatically generated training set. In the second step, a probability model was inferred from the auto-annotated training set. The probabilistic model captures staining patterns in mutually exclusive cell types (CD4^+^, CD8^+^) and builds a single probability model for each marker. Manual annotations of the immune cell types were also used to validate the algorithm performance with accuracy levels ranging from 70-100% for predicted vs annotated cells (150-500 cells annotated per marker, depending on abundance). After cell-level predictions were made for each marker, they were combined to generate multi-marker immune cell classification for each cell.

Analysis involved calculating the distances between immune cells and cancer cells, assessing their proximity to one another (Euclidean distance between the centre points), as the objective of this study was to determine whether spatial proximity was associated with prognosis in stage III CRC patients. We developed a new approach here referred to as region-based nearest neighbourhood analysis. We first clustered the cancer cells within the TMA core and then quantified the intratumoural T cells (T killer cells and T helper cells) within the cancer cell clusters. ‘Hot’ immune clusters were tumour cell clusters containing at least one CD8^+^ T cell and at least one CD4^+^ T cell (containing both T cells). We hypothesised that a higher number of “immune hot” clusters within a core correlated with better outcome and longer recurrence time.

This newly proposed approach consists of three steps: firstly, unsupervised machine learning (ML) was used to cluster tumour regions into smaller dense clusters using k-means clustering with further determination of the optimal number of clusters based on internal validation metrics (elbow method).

Second, we calculated the convex hull of the given set of points. Finally, we used Point-In-Polygon test [22] to count the number at cytotoxic T cells and helper T cells within the cancer cluster.

## Results and Discussion

### The percentage of T-killer cells does not predict recurrence risk in stage III CRC patients

We analysed 462 core samples (number of cells: 1,669,228) from 221 treated stage III patients in our discovery cohort (MSK). The validation cohort, HV, is comprised of 272 samples from 98 patients with stage III CRC. As shown in Figure 2 there was a non-significant trend for patients whose tumours had a high percentage of CD8^+^ T cells to have a better prognosis in the discovery and validation cohort (p=0.074 and p=0.19, respectively). Likewise, the percentage of CD4^+^ T cells in tumours was not significantly associated with disease-free survival (DFS) in both discovery and validation cohorts.

**Figure. 2.**
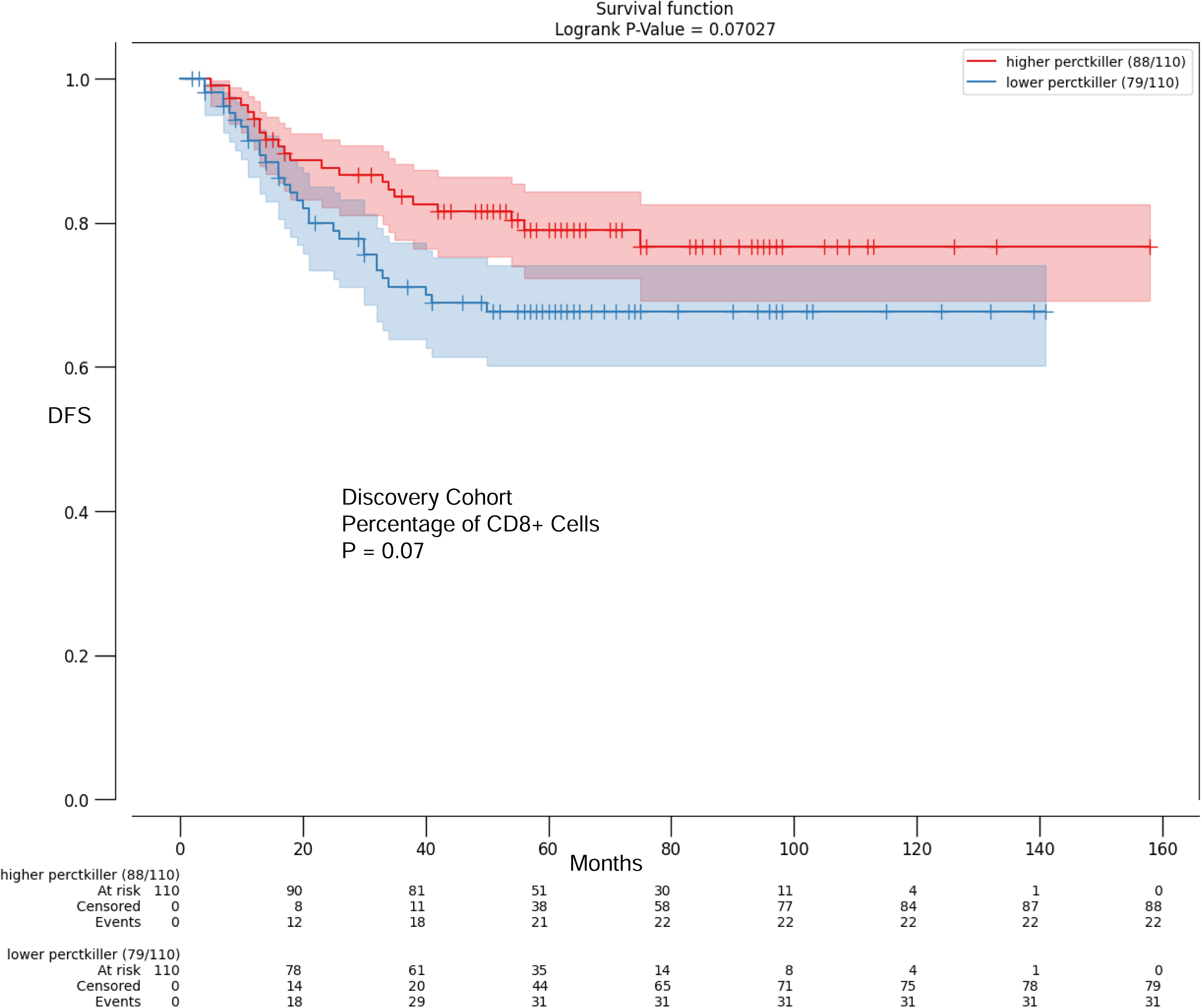
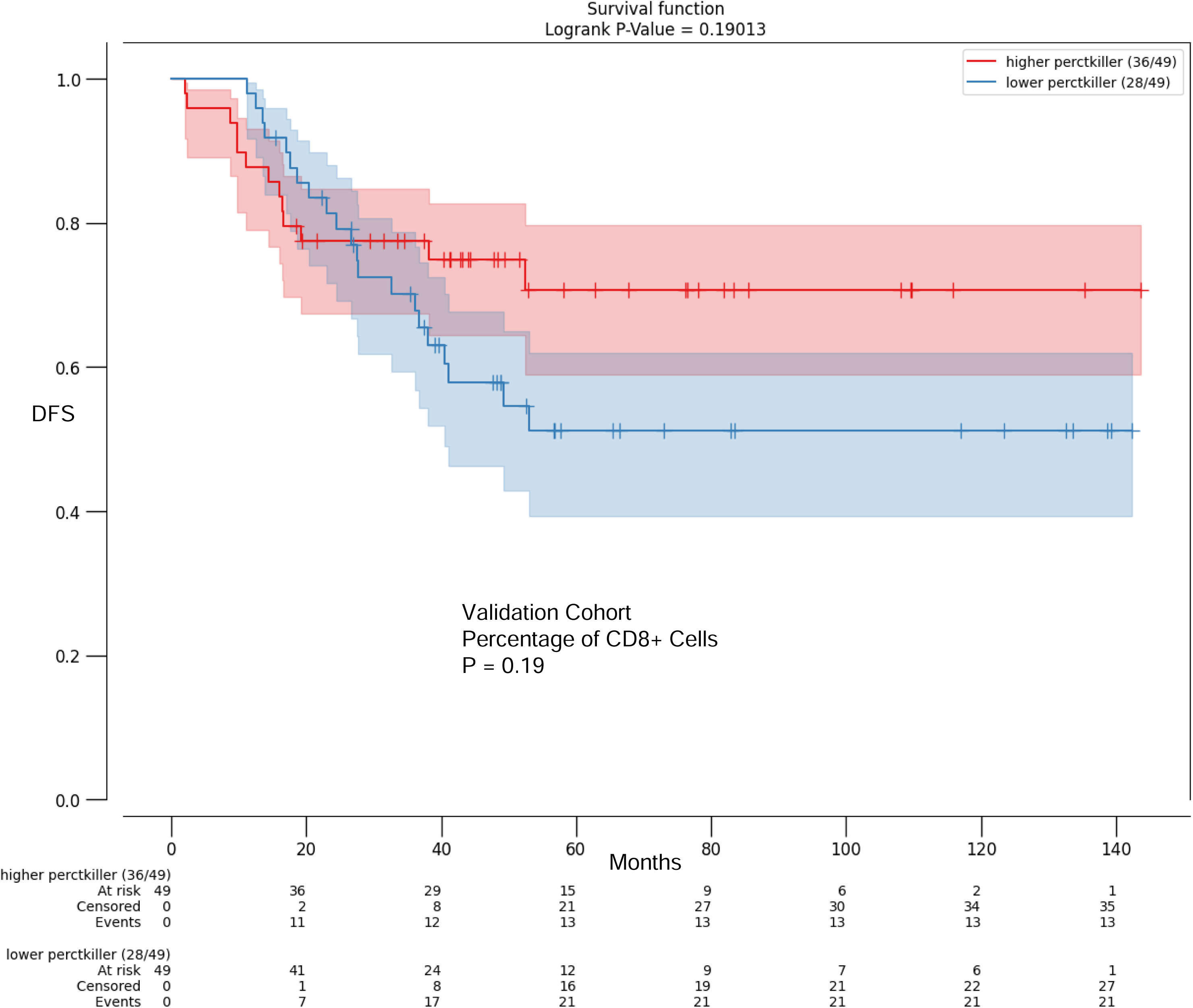
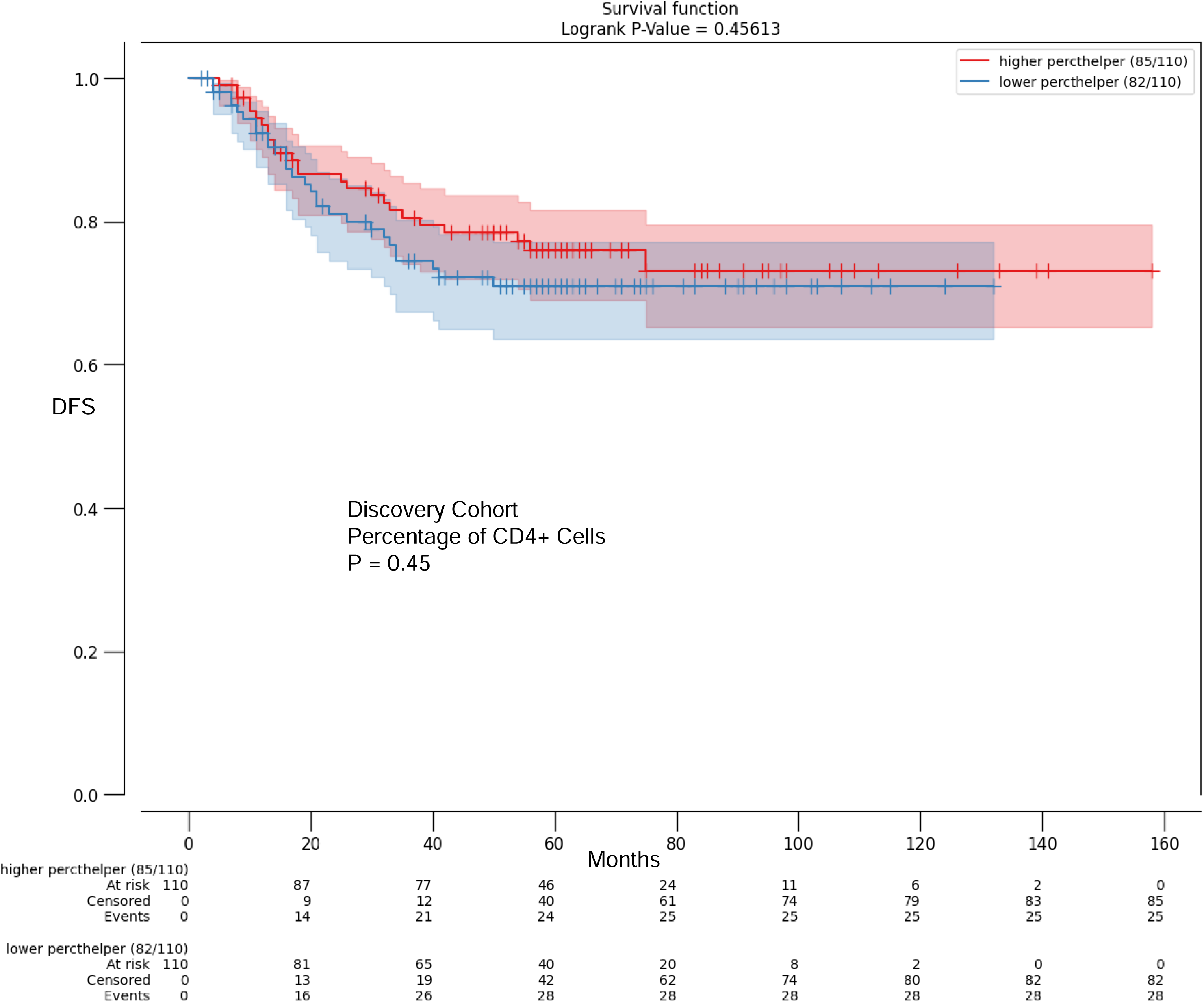
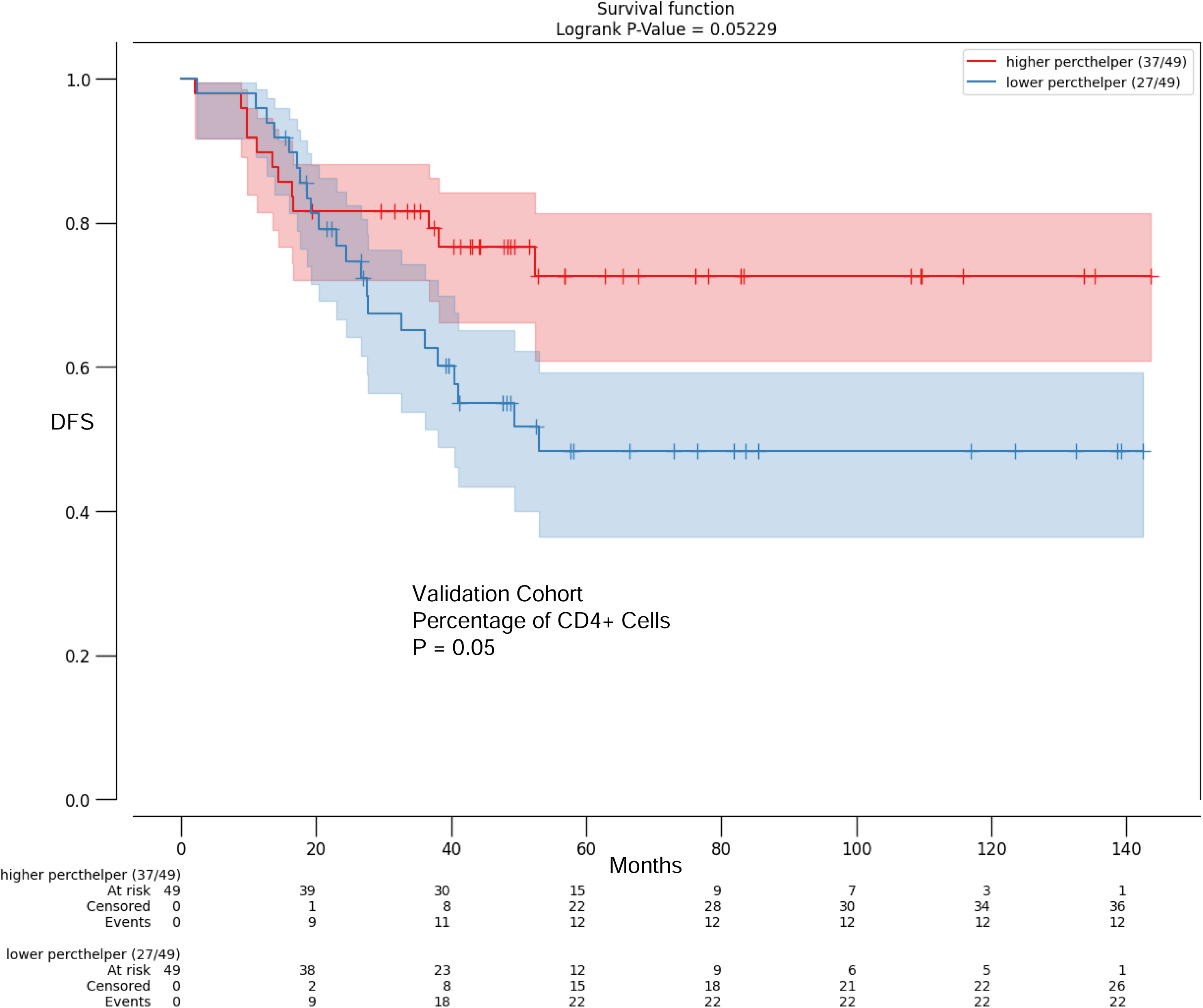
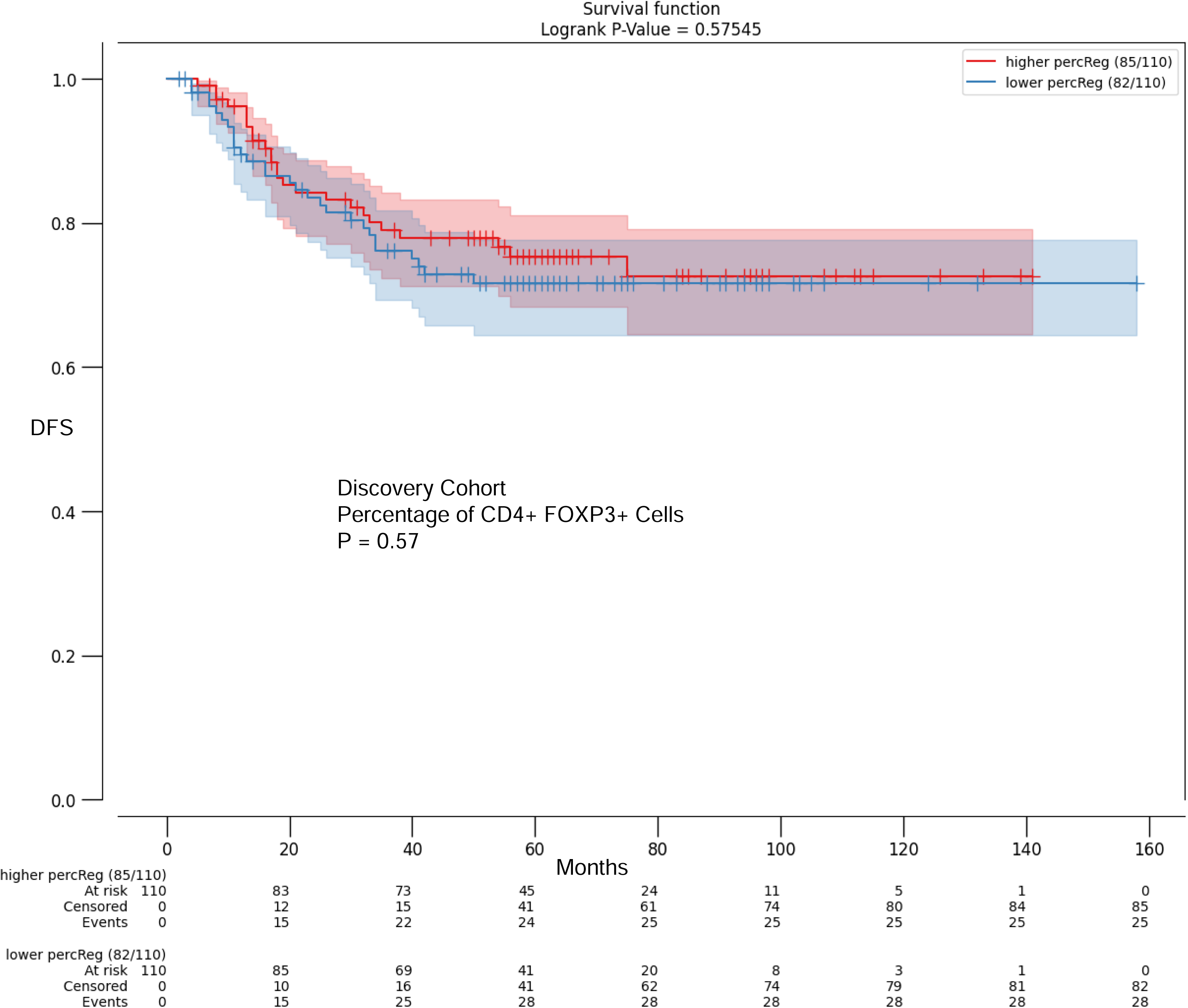
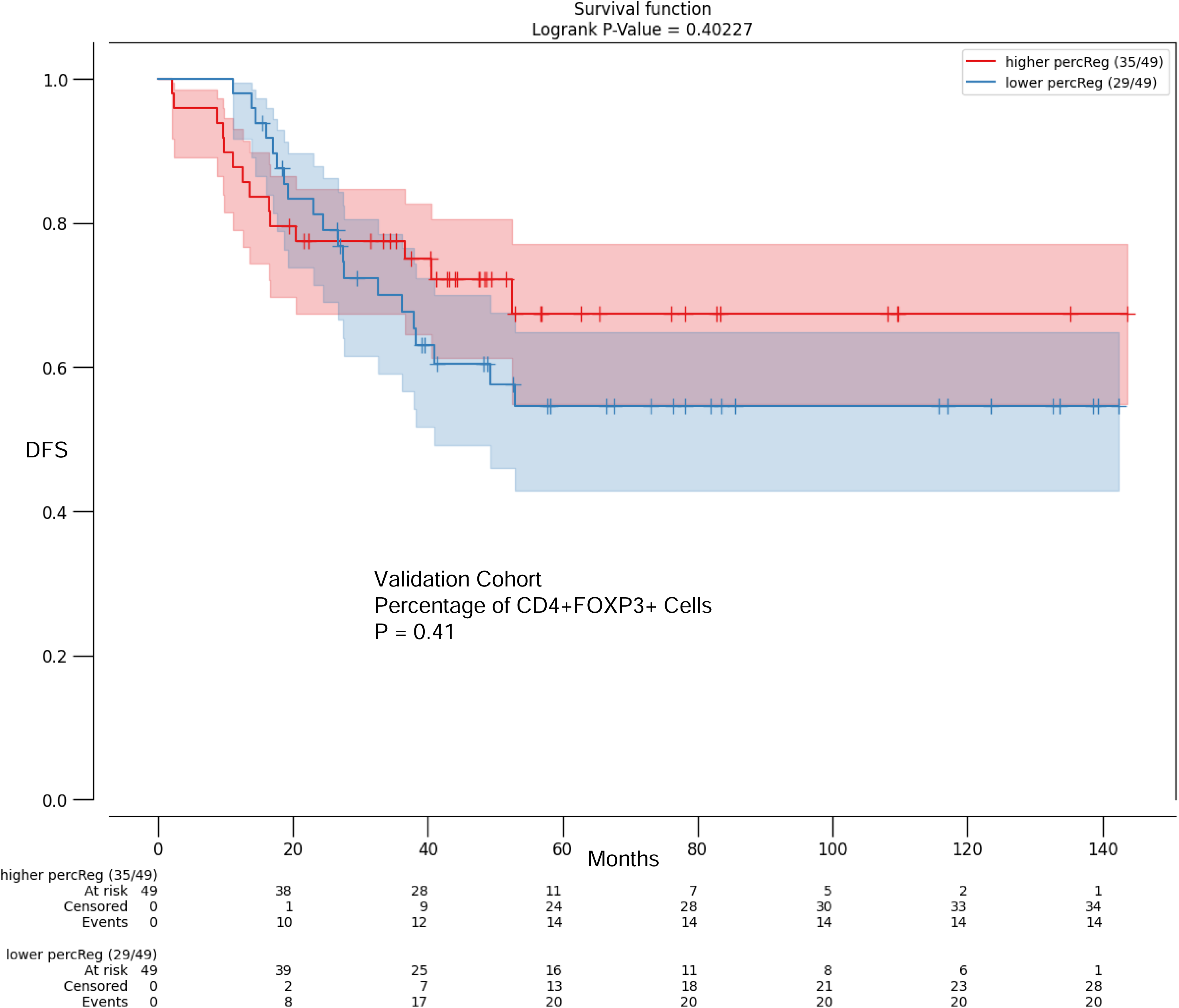
The percentage of Intratumoural T cells does not serve as a prognostic biomarker for the survival of stage III CRC patients treated with 5-FU-based adjuvant chemotherapy. The fraction of CD4^+^ and CD4^+^ FOXP3^+^ T cells did not function as a prognostic indication in the discovery cohort, as shown in Figure. Figures showing the univariate survival analysis for intratumoural T cell densities classified on the median for disease-free survival (DFS) based on Kaplan-Meier curves. The Kaplan-Meier survival curve variations are displayed as a log-rank p value.

### A higher percentage of ‘hot’ tumour clusters is associated with reduced recurrence risk

We hypothesised that the intratumoural T cell spatial distribution within the tumour rather than the absolute numbers of T cells may correlate more closely with patient outcomes. To discover the cell spatial organization and scrutinize T cell–cancer cell interactions, we calculated the percentage of “immune-hot” clusters within a tumour by the use of a region-based nearest neighbourhood analysis approach. The concept of “hot” and “cold” clusters aligns with the idea of immune infiltration impacting tumour cell survival.

As described in Methods, we first clustered cancer cells within a tumour into smaller spatial clusters and then determined whether there was at least one CD8^+^ (T killer) cell in the presents of at least one CD4^+^ T (T helper) cell within each individual spatial cluster thus indicating immune ‘hot’ cluster. The average number of clusters per core was 8.56 ± 1.59 and the average number of cells per cluster was 182.8 ± 73.7.

The percentage of ‘hot’ tumour clusters inside a core (cancer clusters with at least one CD8^+^ T cell and one CD4^+^ T cell) is used as a predictive indicator for the survival of stage III CRC patients. A higher percentage of hot clusters, which contain at least one T killer cell and T helper cell, is connected with a better prognosis (Figure 3). We also compared the number of patients with microsatellite instability among high-risk and low-risk patients. In our discovery cohort, cluster 2 (Low Risk Patients) had more MSI patients, but the difference was not statistically significant (pvalue = 0.15).

**Figure. 3.**
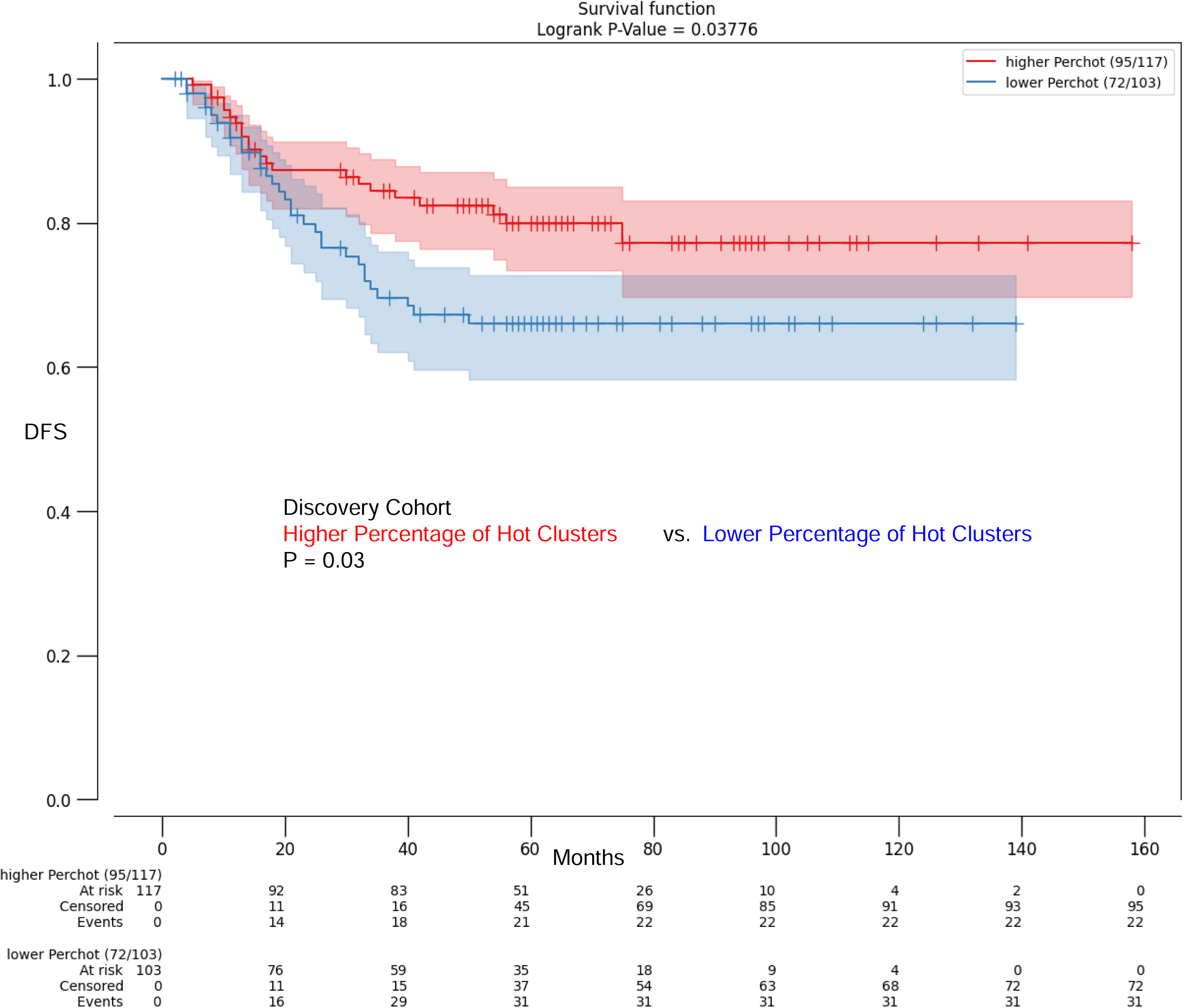
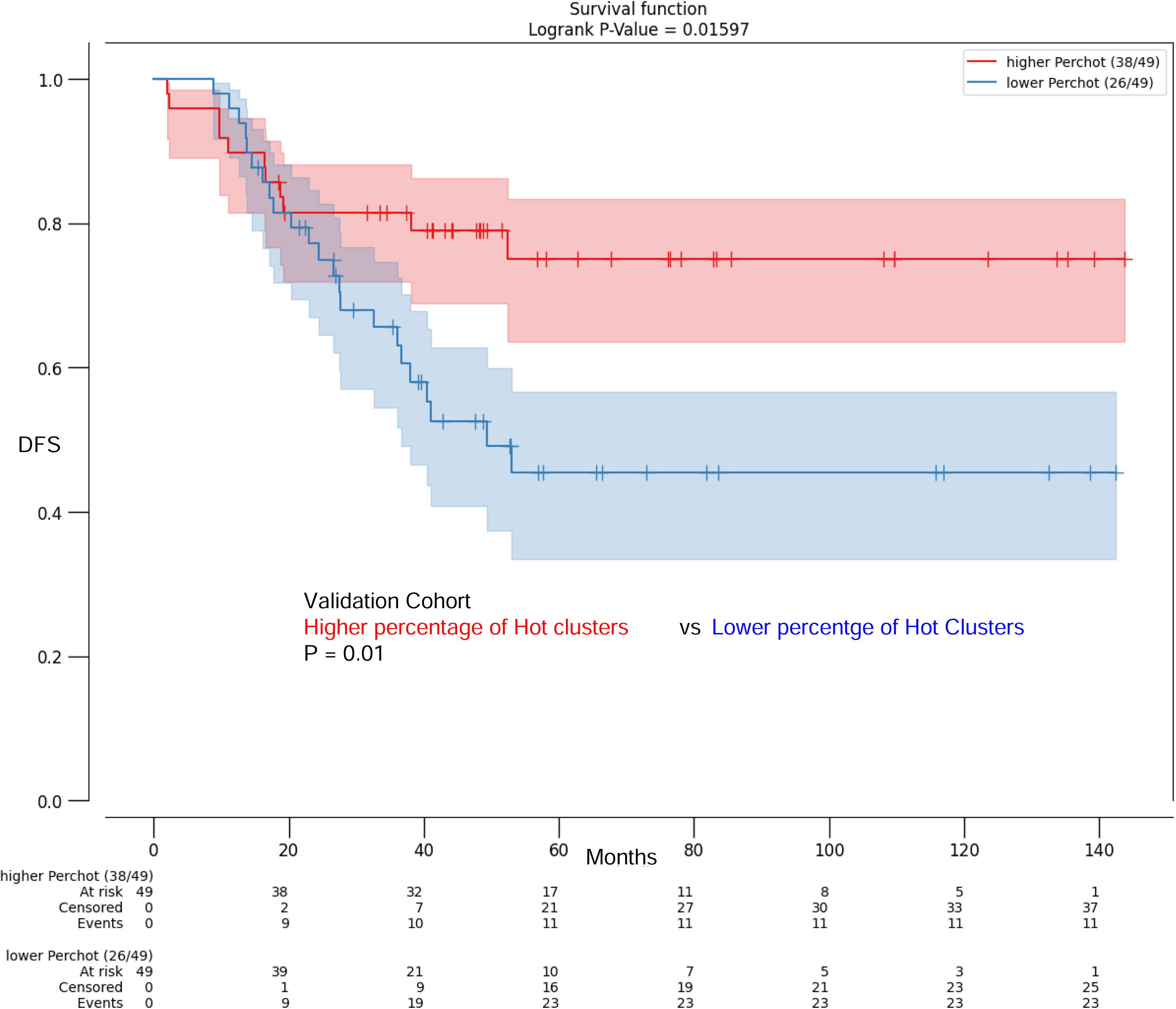
The percentage of ‘hot’ tumour clusters within a core (cancer clusters with at least one CD8^+^ T cell and one CD4^+^ T cell) serves as a prognostic biomarker for the survival of stage iii CRC patients. Higher percentage of the hot clusters - clusters including at least one T killer cell and T helper cell is associated with better prognosis. Figures displaying the categorization of number of hot clusters according to the median disease-free survival (DFS) using Kaplan-Meier curves in the univariate survival analysis. A log-rank p value represents the variability in the Kaplan-Meier survival curve.

### The median distance between CD8^+^ T cells, CD4^+^ T cells and the tumour cells serves as an prognostic biomarker for the DFS of patients after treatments

To build on these findings, we next investigated whether the median distance between cancer cells and intratumoural T cells served as a prognostic factor. Here, we measured the Euclidean distance between cancer cells and their nearest neighbouring T cells (as it is described in Figure 1). The median distance between CD8^+^ T cells and cancer cells in the discovery cohort was 92.77 microns and in the validation cohort 79.22 microns.

As illustrated in Figure 4, the distance between CD8^+^ T cells, CD4^+^ T cells and the tumour cells (Distance between a cancer cell and the nearest CD8^+^ T neighbour cell + Distance between a cancer cell and the nearest CD4^+^ T neighbour cell) served as a strong, prognostic spatial biomarkers in both discovery and validation cohorts (Discovery: p = 0.015, Validation: p = 0.045). Cox proportional hazards models for DFS survival is presented in **Supplementary Table 2**. This table presents P values from multivariable analysis adjusted for the distance between CD8^+^ T cells, CD4^+^ T cells, cancer cells and clinical parameters.

**Figure. 4.**
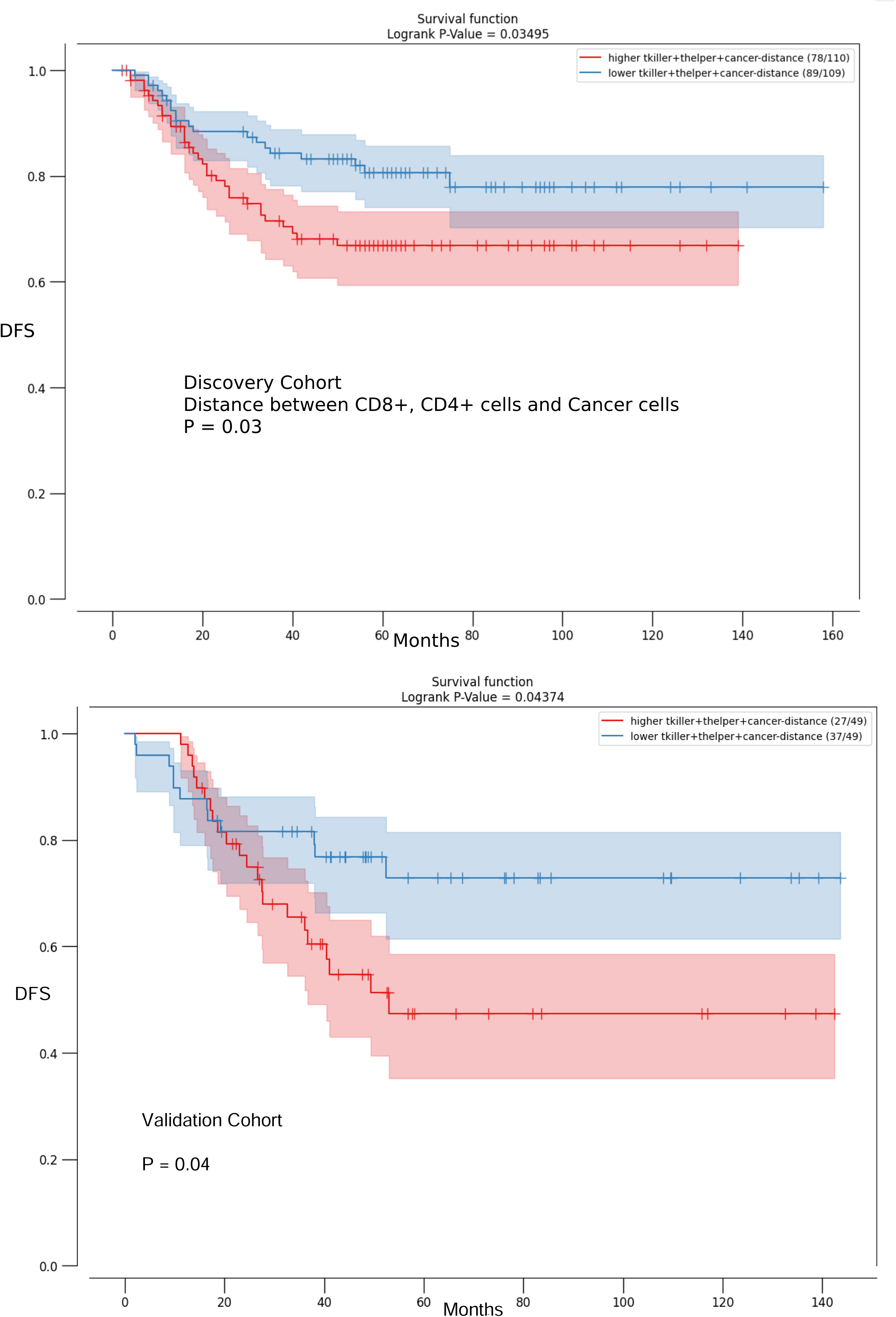
The median distance between CD8^+^ T cells, CD4^+^ T cells and the tumour cells serves as an prognostic biomarker for the survival of patients after treatments. Lower distance between CD8^+^ T cells, CD4^+^ T cells and the tumour cells is associated with better prognosis. Figures depict the categorization of the median distance between CD8^+^ T cells, CD4^+^ T cells, and tumour cells based on median disease-free survival (DFS) using Kaplan-Meier curves in univariate survival analysis. We utilized the log-rank p value to depict the variability in the Kaplan-Meier survival curve.

In contrast, the distance between CD4^+^ T cells and cancer cells was not prognostic (Discovery: p = 0.56, Validation: p = 0.269). The distance between CD4^+^ FOXP3^+^ T cells and cancer cells also showed no prognostic significance based on disease free survival in the discovery cohort (p=0.072). The median distance between CD8^+^ T cells and cancer cells also did not significantly predict DFS (**Supplemental Figure 2**). A statistically significant effect was also observed in the validation cohort for the distance between CD8^+^ T cells, CD4^+^ T cells and the tumour cells.

### Overexpression of Caspase-3, Caspase-8, Ki67 and HLA-1 in cancer cells situated in close proximity to CD8^+^ T cells

Next, we investigated whether the protein signatures of cancer cells situated in close proximity to an intratumoural T cells differed from tumour cells that were not situated in close proximity to an intratumoural T cell. We considered that this may further help develop a refined model that can cluster patients into two low-risk and high-risk groups, based on a combination of both spatial immune and biological biomarkers.

In the discovery cohort (Figure 5) we observed an overexpression of apoptotic regulators (BAK, BCL2, Caspase 3, Caspase 8, Caspase 9) in cancer cells in close proximity to CD8^+^ T cells, suggesting crosstalk between immune cell recognition and apoptosis susceptibility. Overexpression of C-myc, Ki67 and HLA-1 was also observed in cancer cells closer to cytotoxic T cells. These findings were validated for BAX, Caspase 3, Caspase 8, HLA-1, GLUT-1, and Ki67 in the validation cohort.

**Figure. 5.**
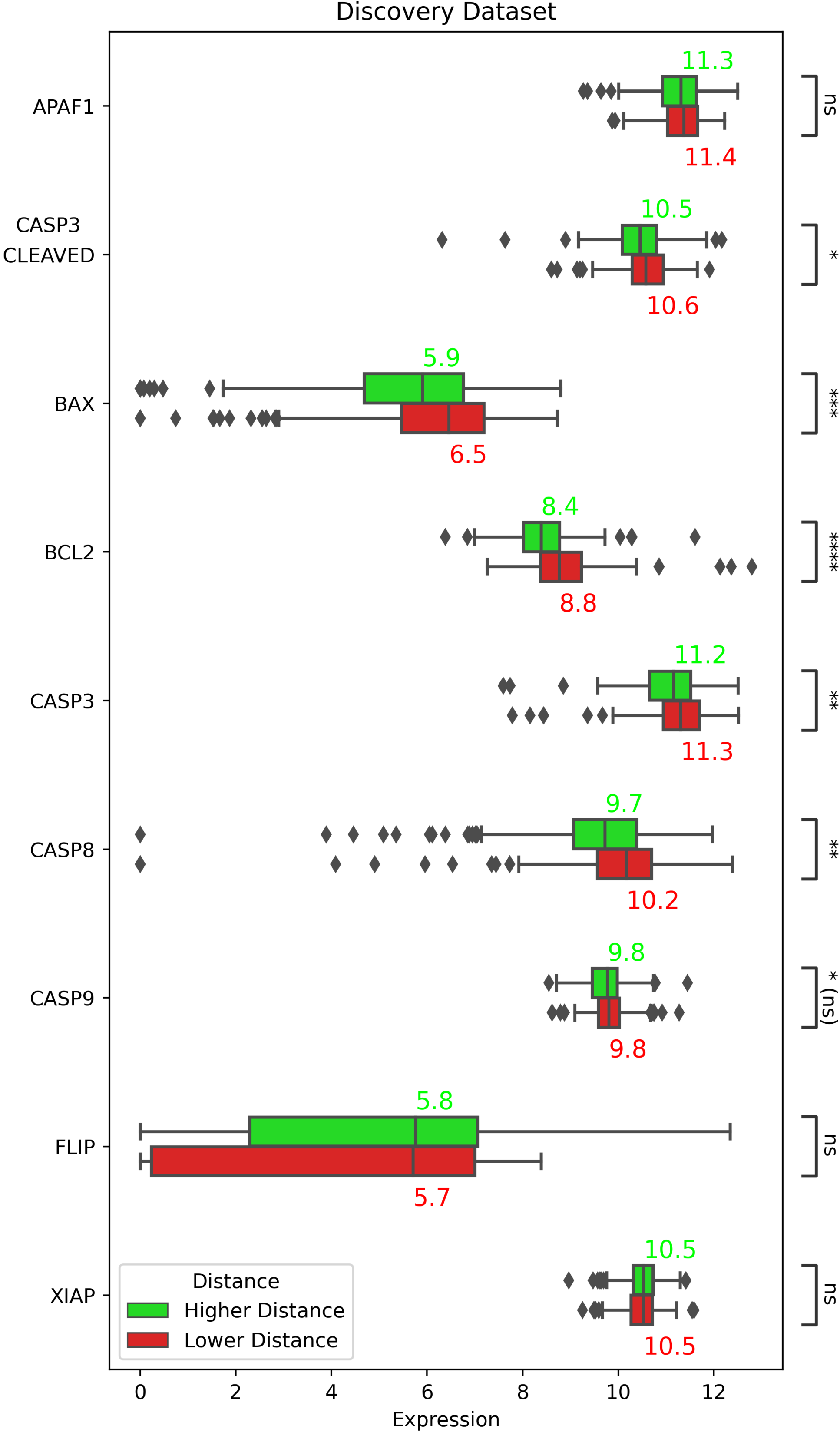
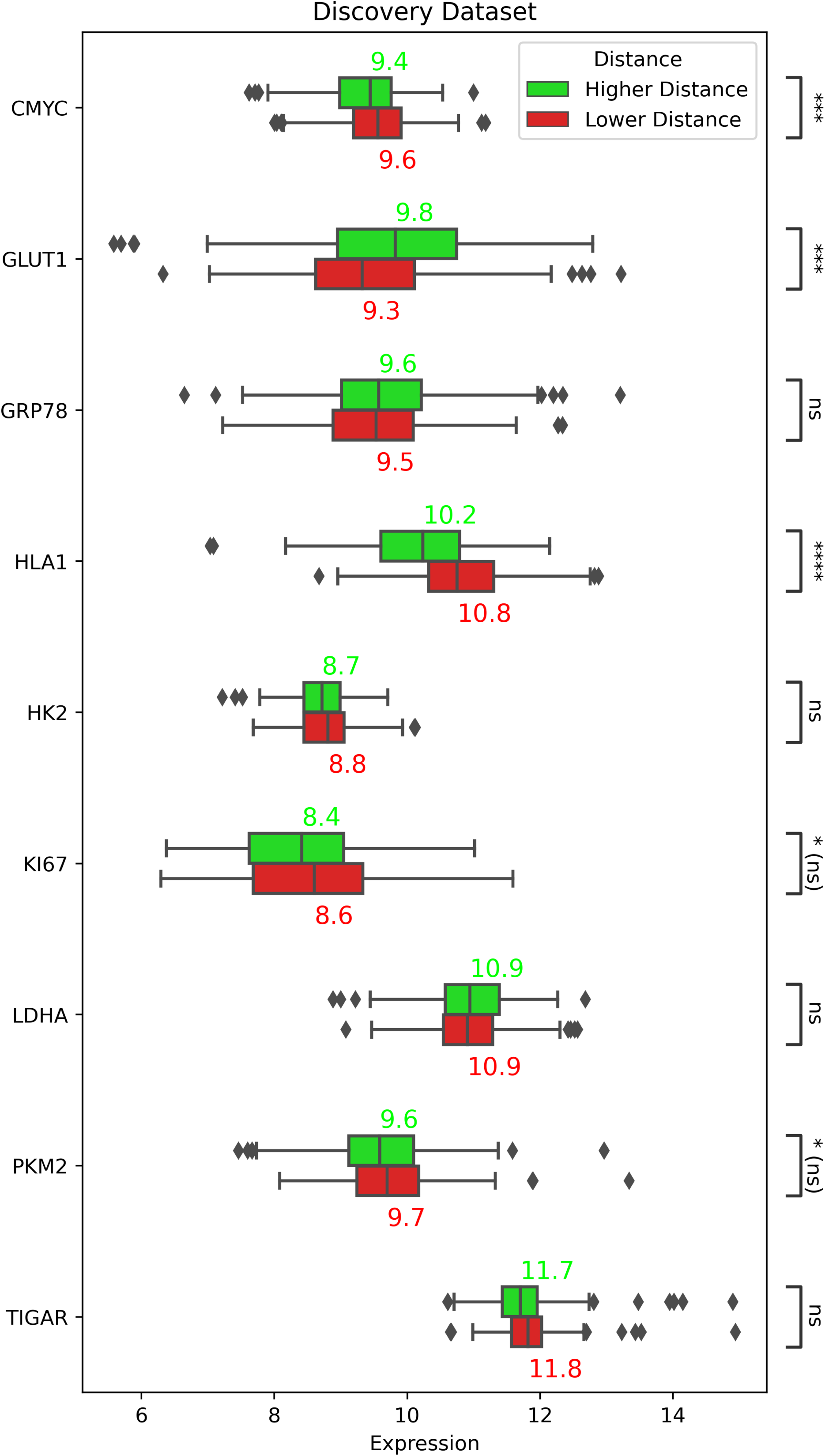
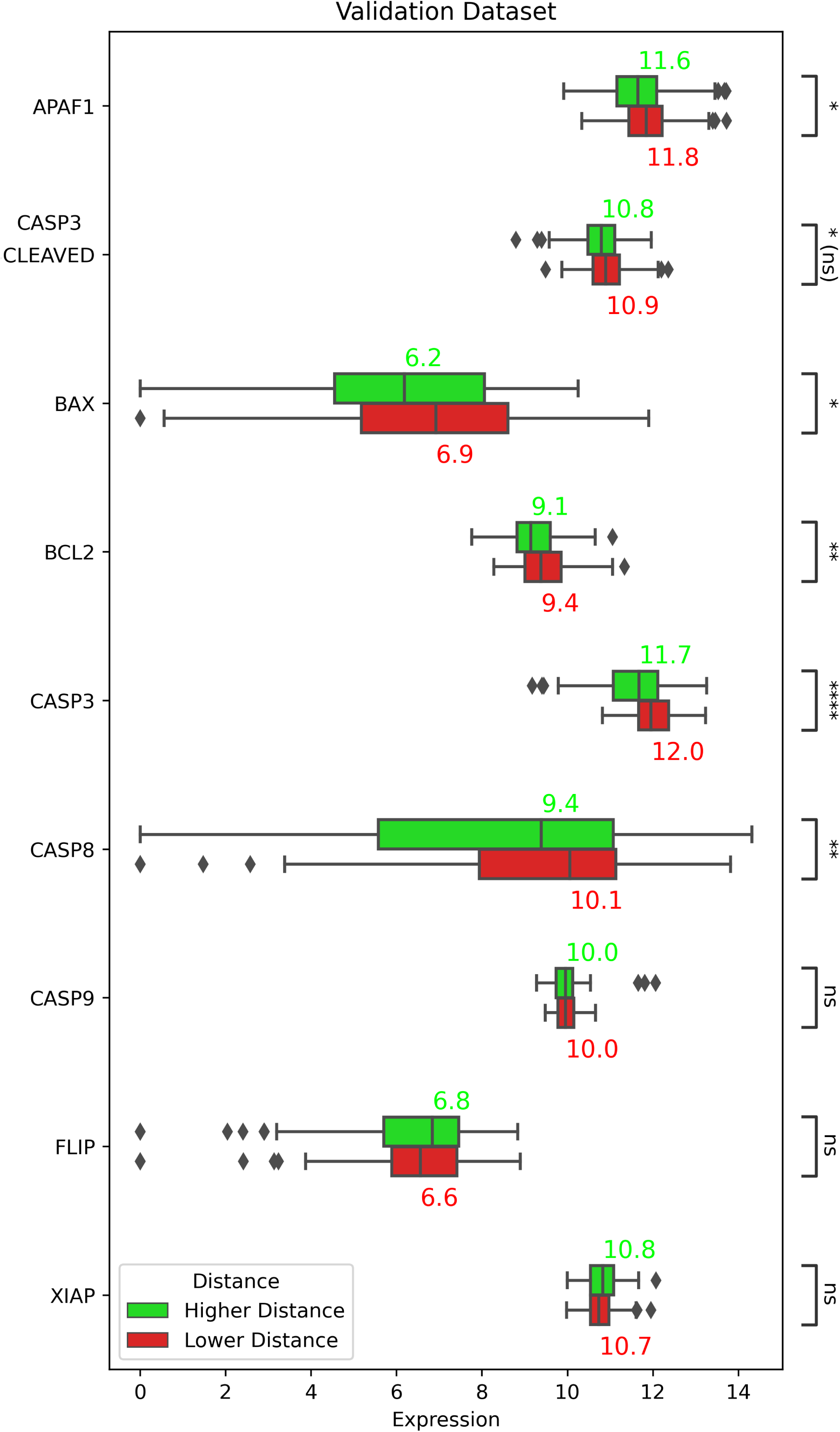
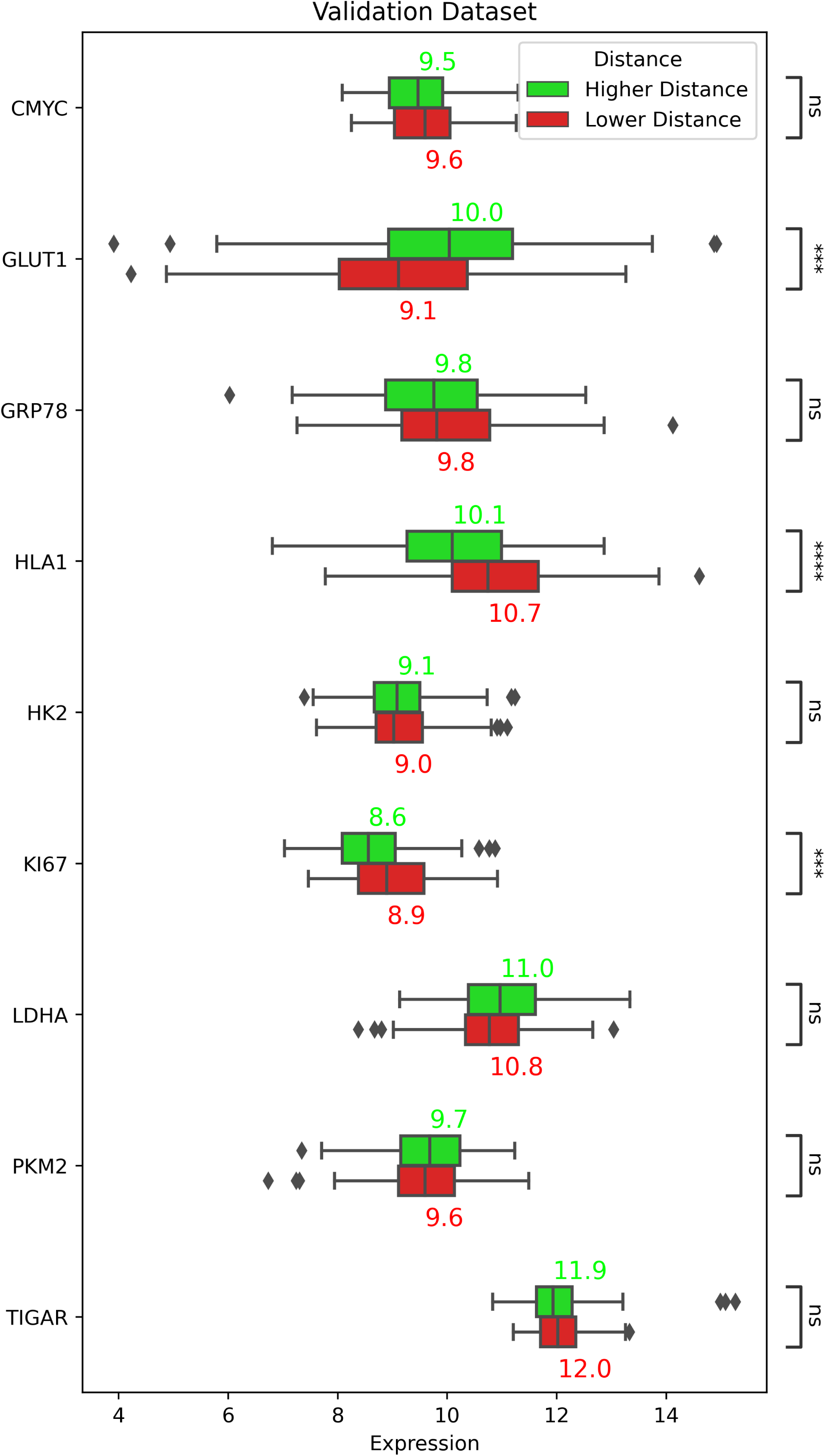
Discovery Dataset/Validation Dataset – Boxplots for cancer hallmarks (Isolated: Cores with higher median distance between cancer cells and CD8^+^ T cells, NonIsolated: Cores with lower median distance between cancer cells and CD8^+^ T cells). Data from the nearest neighbourhood study indicates that cancer cells that are adjacent to CD8^+^ T cells overexpress Caspase-3, cleaved Caspase-3, Caspase-8, Ki67, and HLA-1. Each box represents the distribution of median per core, comparing Isolated (higher than median distance between cancer cells and CD8^+^ T cells) and Non Isolated (lower than median distance between cancer cells and CD8^+^ T cells) groups.

We also detected altered levels of APAF1 in proximity to T Helper cells (**Supplemental Figure 3**) and altered levels of BCL2 in proximity to CD4^+^ FOXP3^+^ T cells (**Supplemental Figure 4**).

### Two distinct clusters within stage III colorectal cancer tumours, with significant differences in HLA-1, Ki67, C-myc, GLUT1, Caspase3, and immune cells spatial features are highly prognostic in stage III CRC

The relationship between cancer hallmarks expression, intratumoural T cell infiltration, and cancer recurrence is complex and may vary depending on many different parameters in the cancer microenvironment. Based on protein signature data and nearest neighbourhood analysis results, we used unsupervised clustering to perform an in-depth investigation of the tumours. We discovered two separate groups based on several pre-investigated factors such as the median distances between immune cells and cancer cells and protein expression patterns. We identified two key spatial biomarkers: the median distance between T Killer cells and cancer cells, and the median distance between T Helper Regulatory cells and cancer cells (**Supplemental Figure 5**). In addition to spatial markers, we next included five biological biomarkers based on our above results (expression of Caspase 3, Ki67, C-myc, GLUT-1, and HLA-1 in cancer cells). The color-coded z-score matrix, is shown in Figure 6.

**Figure. 6.**
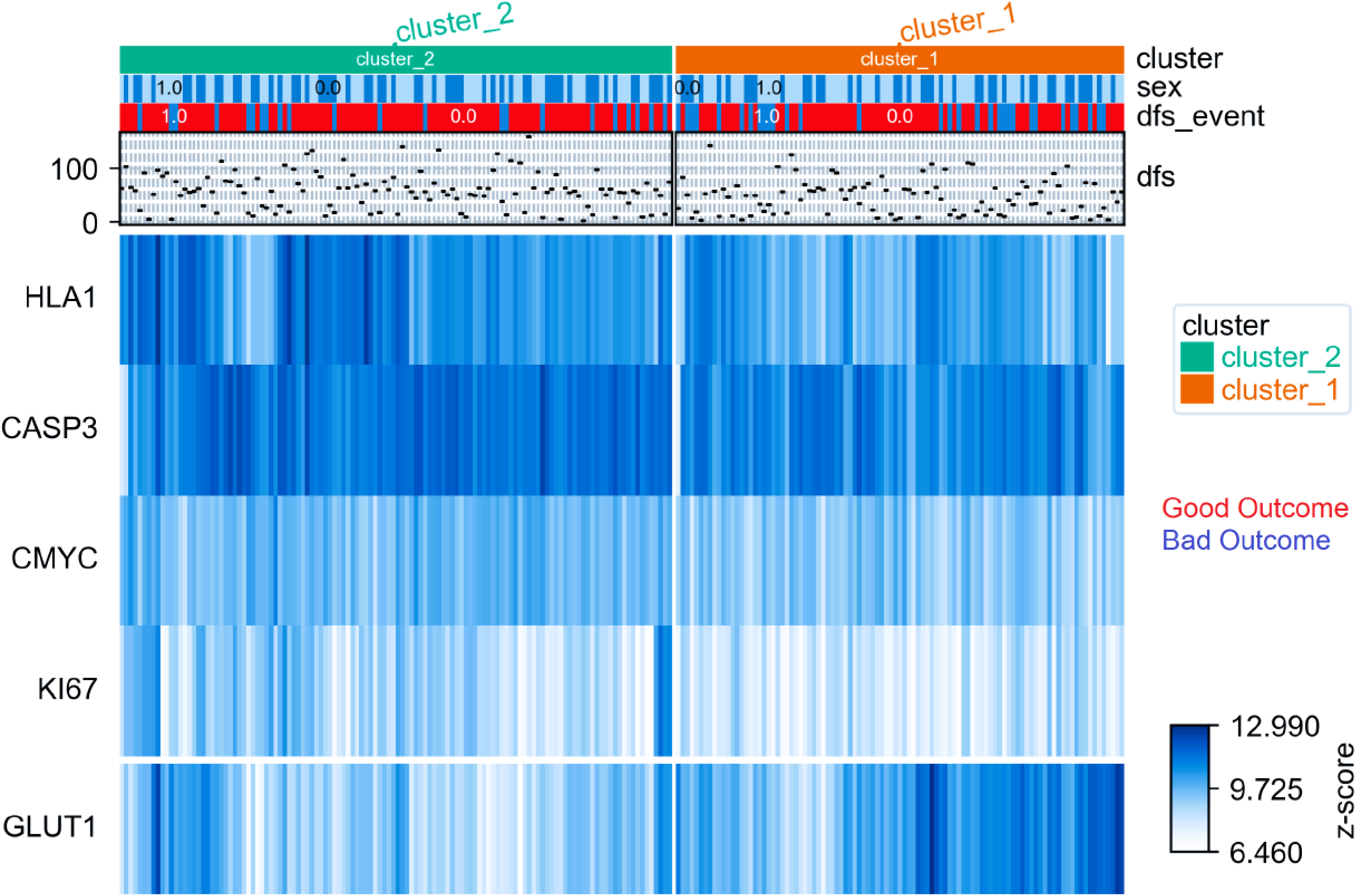
ClusterMap – Discovery Cohort : Expression of Caspase 3, Ki67, C-myc, GLUT-1, and HLA-1 in cancer cells, the median distance between T Killer cells and cancer cells, and the median distance between T Helper Regulatory cells and cancer cells. These values quantify the average distance between cancer cells and cytotoxic/regulatory T cells, illuminating the spatial interactions between them and the immunological environment of the tumour. We have taken into consideration five biological indicators (expression of Caspase-3, Ki67, C-myc, GLUT-1, and HLA-1 in cancer cells) in addition to spatial markers. The blue-white color-code displays z-score.

The observation of high expression of HLA-1 and Caspase 3 in cluster 2, along with low expression of GLUT1 in cancer cells in this cluster, suggests important differences in the tumour microenvironment between the two clusters. We observed low expression of HLA-1, C-myc, Caspase 3 and Ki67 in tumour cancers within cores belong to cluster 1 (**Supplemental Figure 5**). We observed significant survival difference among stage III CRC patients (Figure 7), demonstrating that an approach combining spatial and biological data delivers the most accurate prognosis (discovery: p-value = 0.004 and validation: p-value = 0.003).

**Figure. 7.**
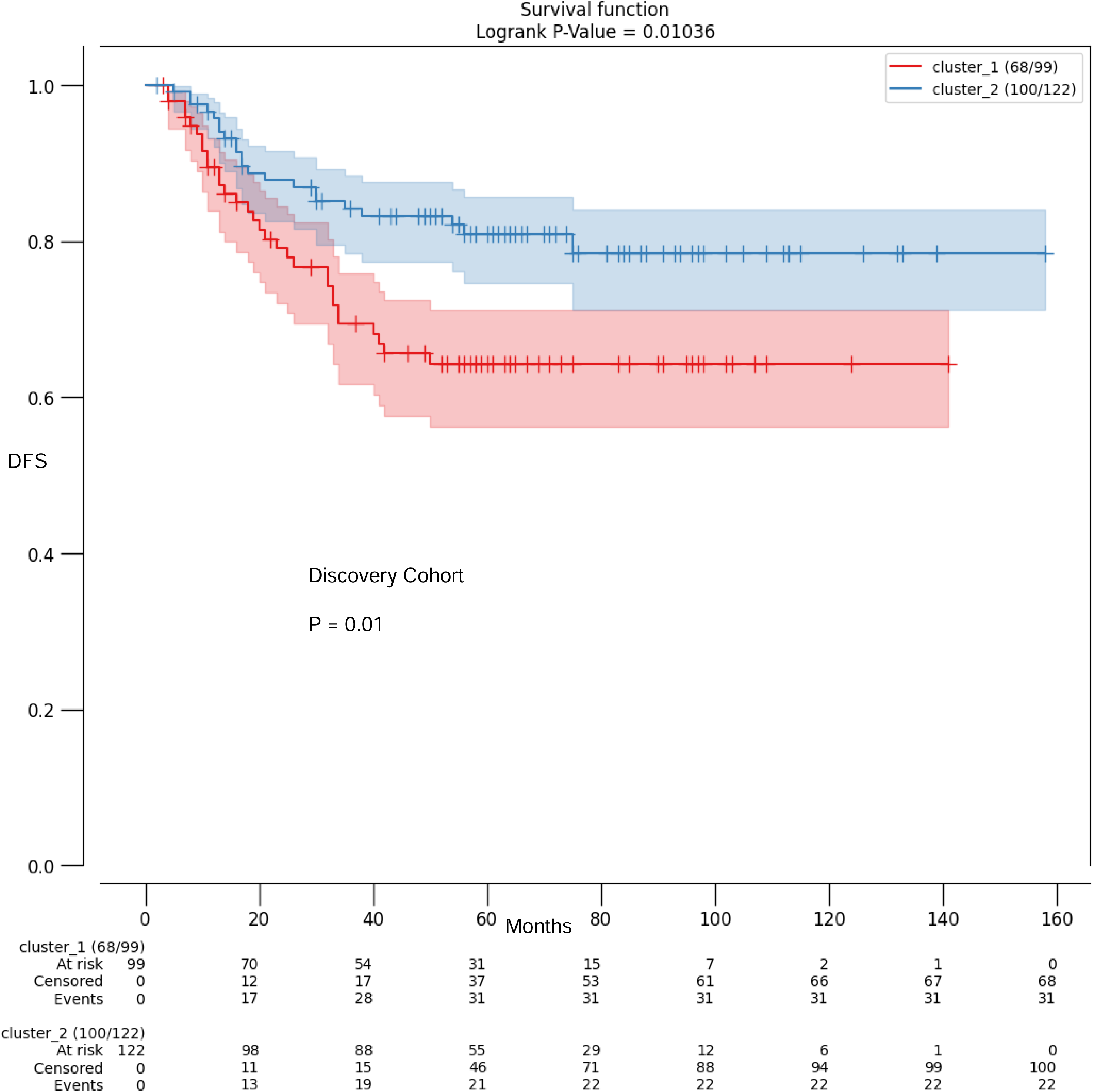
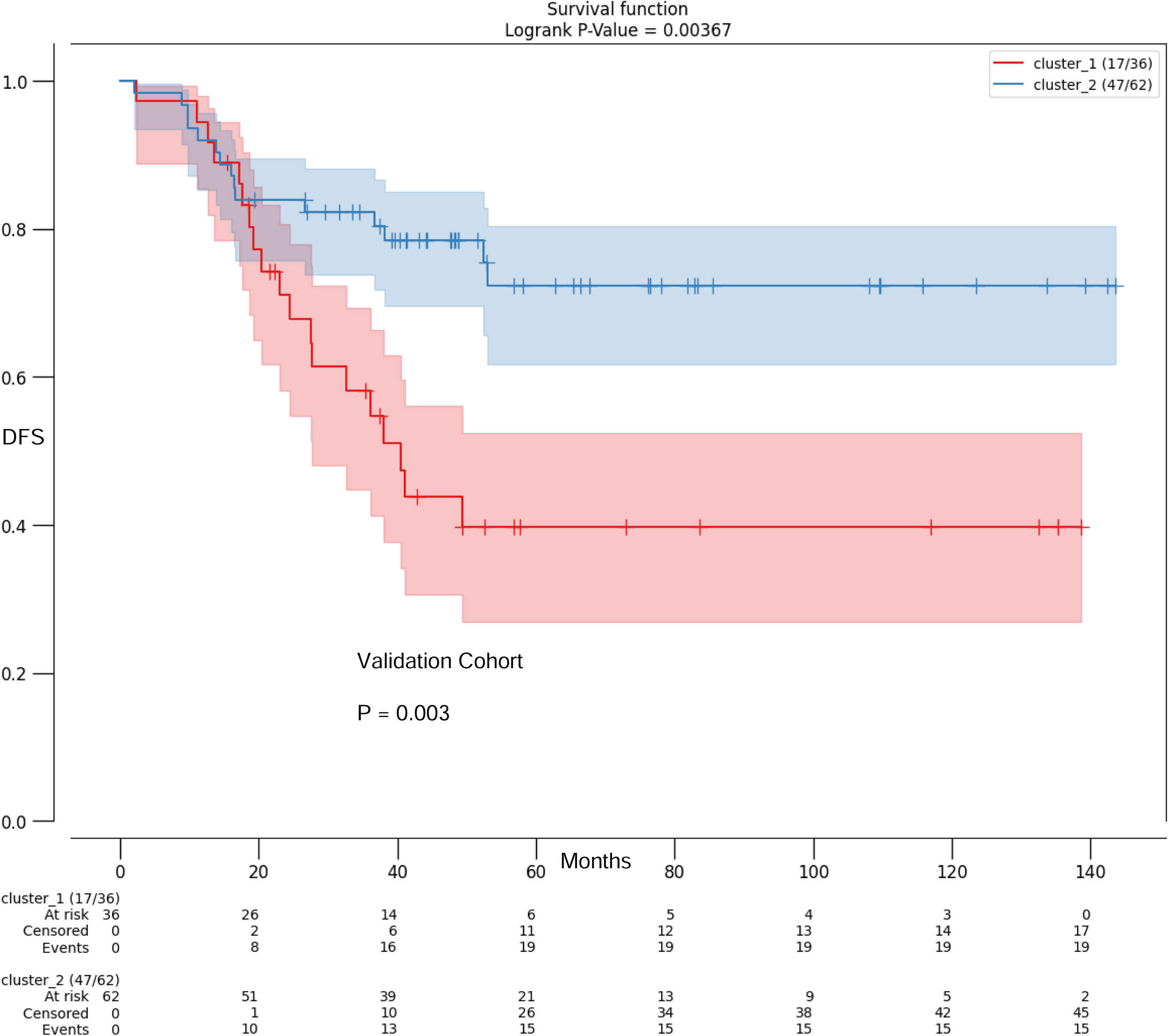
The considerable survival difference among patients with cores in cluster two is a positive finding, indicating that these clusters may have therapeutic importance in predicting patient outcomes (in both the discovery and validation groups). Kaplan-Meier plots illustrating DFS for two distinct risk groups: Cluster 1, high risk patients and Cluster 2, low risk patients. Noteworthy divergence in survival curves highlights the prognostic significance of risk-based clustering. A log-rank p value represents the variability in the Kaplan-Meier survival curve.

In the present study we showed that the distance between CD8^+^ T cells, CD4^+^ T cells and the tumour cells (using a nearest neighbourhood approach) serve as prognostic spatial biomarkers in stage III CRC. We also provide evidence that the percentage of intratumoural T cells within tumours may not necessarily be considered as a prognostic biomarker. The findings of this study also suggest that there are two distinct clusters within stage III CRC tumours, with significant differences in expression of cancer hallmarks and immune cells spatial organization between them. Furthermore, we have noted that these clusters exhibit improved prognostic power, indicating significant clinical relevance.

We investigated the spatial arrangement of CD8^+^ T cells, CD4^+^ T cells and cancer cells within the tumour microenvironment in stage III CRC patients. Previous reports [23] utilized spatial image analysis approaches to characterize tumour-immune interactions. Wang et al [23] showed that the fractions of proliferating CD8^+^ T cells and MHCII^+^ cancer cells were dominant predictors of response, followed by cancer–immune interactions with B cells and granzyme B^+^ T cells.

To the best of our knowledge, this is the first study utilising nearest neighbour approach and region-based nearest neighbour approach for analysing cancer cell - T cell interactions in hyperplexed stage III CRC tissues. Through comprehensive analysis and validation, we reveal that the distance between CD8^+^ T cells, CD4^+^ T cells and cancer cells is a robust predictor of patient prognosis. The results of nearest neighbourhood analysis can provide valuable information about the degree of immune infiltration, cell clustering, and functional associations within the tumour microenvironment. Highplexing techniques are powerful tools that allow us to simultaneously analyse spatial patterns but also the expression of multiple biomarkers in individual cancer cells.

The notion of using derivatives of nearest neighbour methods and cell proportion quantification for identifying clusters has been tackled also in previous literature [24, 25]. In our study, each tumor was sampled in triplicate from the centre of the tumour and the single cell data is not extracted from luminal side and advancing edge [26].

Our results reveal that while there were trends for intratumoural T cell infiltration and reduced recurrence risk across all the CRC cohorts, they were not significant and the percentage of cancer clusters with CD8^+^ T cells and CD4^+^ T cells inside (using region-based nearest neighbourhood approach) served as prognostic spatial biomarkers in stage III CRC.

The findings of the latest part of this study suggest that there are two distinct clusters within stage III colorectal cancer tumours, with significant differences in these spatial and biological biomarkers between them. The technique is based on a combination of spatial and biological biomarkers, enhancing our understanding of the tumour microenvironment and its relevance to patient outcomes. Cluster 1’s underexpression of GLUT-1 and overexpression of HLA-1 and caspase-3 in cancer cells point to significant changes in the tumour microenvironment between cluster 1 and cluster 2. Furthermore, we demonstrated the potential therapeutic importance of these clusters in predicting patient outcomes. The overexpression of Caspase-3 in cancer cells closer to CD8^+^ T cells can be considered as biological evidence that indicates that the immune system, specifically CD8^+^ T cells, is actively engaging with and targeting cancer cells for destruction. The overexpression of HLA-1 (Human Leukocyte Antigen 1) molecules on cancer cells can potentially lead to a better outcome for cancer patients. In locally advanced colorectal cancer, a high expression of HLA-1 is linked to a better tumour prognosis. HLA-1 molecules play a crucial role in presenting antigens to immune cells, such as T cells [18]. Our data from the nearest neighbourhood study reveals that cancer cells that were adjacent to intratumoural T cells overexpressed HLA-1. In our cohort of stage III patients treated with adjuvant chemotherapy, the overexpression of HLA-1 on cancer cells could potentially also enhance the immune response triggered by the treatment.

We also demonstrate that cancer cells that are close to T cells overexpress Ki67. Ki67 is a marker for cellular proliferation within a tumour. The overexpression of the protein Ki67 in cancer cells located in proximity to CD8^+^ T cells could be indicative of an active immune response specifically against proliferating cancer cells which may provide additional cues for recognition to immune cells. The increased expression of Ki67 in cancer cells near CD8^+^ T cells may also be a response to the immune system’s attempt to eliminate the cancer. Interestingly a previous study [27] reported that elevated Ki67 levels were significantly associated with a better outcome in stage III CRC patients, but only in patients who received adjuvant chemotherapy (P=0.007). In this context, we observed higher Ki67 levels in cancer cells is proximity to CD8^+^ T cells in cluster ‘1’ which was associated with an improved disease-free survival after adjuvant chemotherapy. The under expression of GLUT1 (Glucose Transporter 1) in cancer cells that were close to CD8^+^ cells, can have several potential implications. CD8^+^ T cells are highly metabolically active, and they require glucose to perform their effector functions, such as killing cancer cells. If cancer cells near CD8^+^ cells under express GLUT1, they may take up less glucose, which could potentially create a metabolic advantage for CD8^+^ T cells and enhance their activity. In line with this hypothesis, a previous study demonstrated that positive GLUT-1 staining after chemo radiation therapy [28] was linked to a poorer outcome. Additionally, lower glucose availability in cancer cells could lead to reduced glycolytic activity within the tumour microenvironment. This might create an environment that is less supportive of cancer cell survival and growth. It should also be noted that our study focused on tumour cores only, and that events occurring at tumour margins may have been missed.

## Conclusions

We here developed a new methodology and revealed new spatial/biological biomarkers for the prognosis of adjuvant chemotherapy treated stage III CRC patients. It underscores the importance of understanding the tumour microenvironment and its implications for patient outcomes in CRC. The median distance between CD8^+^ T cells, CD4^+^ T cells and cancer cells and the formation of ‘hot’ clusters, rather than the proportion of intratumoural T cells, functioned as prognostic biomarker for patient survival following therapy. We also show that cancer cells in close proximity to CD8+ T cells overexpress Caspase-3, Caspase-8, Ki67, and HLA-1. Integration of these signature provides novel and improved prognostic biomarkers for stage III CRC.

## Funding

Research reported in this publication was supported by the National Cancer Institute of the National Institutes of Health under award number R01CA208179 (PI Ginty). DBL was supported by a US-Ireland Tripartite R01 award (NI Partner supported by HSCNI, STL/5715/15). JHMP is supported by US-Ireland Tripartite award from Science Foundation Ireland and the Health Research Board (16/US/3301). JS is supported by R01CA208179 and NCI grant NCI-P30 CA 008748.

## Data availability

Imaging data, cell masks, and generated single-cell measurements of 56 markers are available from the lead contact. Further information and request for code or resources should be directed to and will be fulfilled by the lead contact.

## The Immunofluorescence imaging of patient TMAs

Please see ref.[21]

## Image pre-processing and quality control

Please see ref.[21]

## Post pre-processing and batch correction

Please see ref.[21]

## Antibody validation and conjugation

Please see ref.[21]

## Appendix.1. Method for counting number of T cells within cancer cell clusters

We used K-means clustering technique and processed cancer cells spatial features (X and Y centroids) to group cancer cells into smaller clusters. The optimal number of clusters within a tumour is calculated by the use of elbow method. The cancer zones were found based on the position of cancer cells within the determined clusters. The convex hull is the smallest convex polygon that encloses all the points. We used the Graham scan algorithm:

- Finding the lowest point: Consider a cluster of cancer cells as set of points P with n points. To find the lowest point:
- Sorting by the polar angle: Given the lowest point, sorting the remaining points based on their polar angles with respect to this point
- Scanning through points: checking whether a new point forms a left turn or a right turn with the last two points in the convex hull

The output of sign operator is positive for counter-clockwise turns and negative for clockwise turns.

Finally, after achieving the convex hulls (the closed curves or the cancer cell clusters), we used Point-In-Polygon test [22] to count the number at cytotoxic T cells and helper T cells within the cancer cluster.

## Supplementary Material

***Supplemental Figure 1.** QC workflow*

***Supplemental Figure 2.** The median distance between T cells and the tumour cells does not serve as prognostic biomarker for the survival of patients after treatments. Figures show how Kaplan-Meier curves from univariate survival analysis are used to classify the median distance between intratumoural T cells and cancer cells based on median disease-free survival (DFS). The Kaplan-Meier survival curve’s variability was represented by the log-rank p value.*

***Supplemental Figure 3.** Discovery Dataset/Validation Dataset – Boxplots for the cancer hallmarks (Isolated: Cores with higher median distance between cancer cells and CD4^+^ T cells, NonIsolated: Cores with lower median distance between cancer cells and CD4^+^ T cells). Comparing the Isolated and Non-Isolated groups, each box shows the distribution of median per core.*

***Supplemental Figure 4.** Discovery Dataset/Validation Dataset – Boxplots for the cancer hallmarks (Isolated: Cores with higher median distance between cancer cells and CD4^+^ FOXP3^+^ T cells, NonIsolated: Cores with lower median distance between cancer cells and CD4^+^ FOXP3^+^ T cells). Each box displays the median distribution for each core when comparing the isolated and non-isolated groups.*

***Supplemental Figure 5.** The observation of overexpression of HLA1 and CASP3 in cluster 2, along with underexpression of GLUT1 in cancer cells in that cluster, suggests important differences in the tumour microenvironment between the two clusters. a-Each box represents the distribution of median per core, comparing High risk patients (Cluster 1) and Low risk patients (Cluster 2) groups. b-The scatter plots for the patient clusters at low and high risk are displayed in the figure. One median per subject is shown by each dot. c-The dendrogram produced by hierarchical clustering using the average linkage between the validation and discovery datasets is displayed in this figure. In addition to spatial biomarkers, we have selected five biological indicators: expression of caspase 3, Ki67, C-Myc, Glut1, and HLA1 in cancer cells.*

**Supplemental Table 1.**
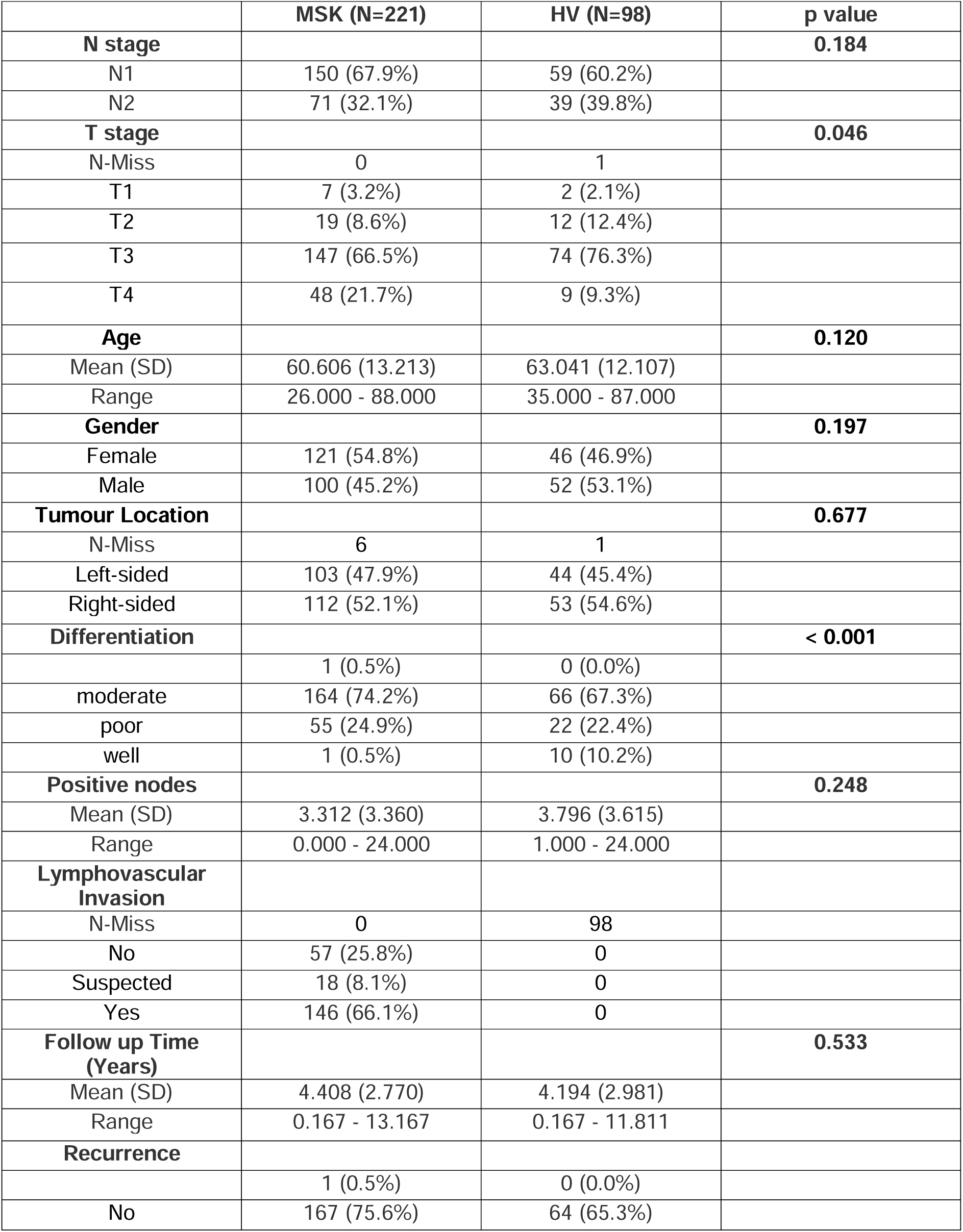
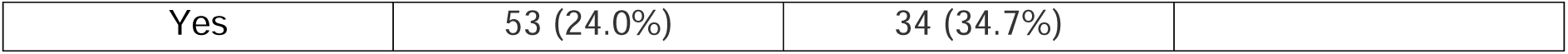
Summary of demographics of the cohorts.

**Supplementary Table 2.** Cox proportional hazards models for DFS survival. P values from multivariable analysis adjusted for Distance between CD4+, CD8+ and cancer cells, age, gender, differentiation and lymphovascular invasion status.

|  | N<br>(Discovery<br>cohort) | P - value | Univariate<br>HR (95% CI) | P<br>value | Multivariate<br>HR (95%CI) |
| --- | --- | --- | --- | --- | --- |
| **Age** |  | **0.71** | **1.13** |  |  |
| Less than 50 | 53 |  |  |  |  |
| More than 50 | 167 |  |  |  |  |
| **Gender** |  | **0.61** | **0.87** |  |  |
| Male | 100 |  |  |  |  |
| Female | 120 |  |  |  |  |
| **Differentiation** |  | **0.66** | **0.87** |  |  |
| Moderate | 163 |  |  |  |  |
| Not moderate | 57 |  |  |  |  |
| **has_lymphovascular_invasion** |  | **0.09** | **1.75** |  |  |
| Yes | 145 |  |  |  |  |
| Other | 75 |  |  |  |  |
| **D between t helper, T killer**<br>**and cancer cells** |  | **0.04** | **1.81** | **0.03** | **1.83** |
| High | 110 |  |  |  |  |
| Low | 110 |  |  |  |  |

## Notes

No conflicts of interest were declared.

### Competing Interest Statement

The authors have declared no competing interest.

### Summary of Updates

Further information and request for code or resources should be directed to by the lead contact. This information has now been added to the manuscript. We have also moved the detailed description of methods to our supplementary data (appendix 1) and have replaced the main text with a simpler description

## References

[1] Hou, Wanting, Cheng Yi, and Hong Zhu. “Predictive biomarkers of colon cancer immunotherapy: Present and future.” Frontiers in Immunology 13 (2022): 1032314.

[2] Killock, David. “Early MRD predicts disease recurrence and benefit from adjuvant chemotherapy in CRC.” Nature Reviews Clinical Oncology 20.3 (2023): 137–137.

[3] Tomasello, Gianluca, et al. “Survival benefit with adjuvant chemotherapy in stage III microsatellite-high/deficient mismatch repair colon cancer: a systematic review and meta-analysis.” Scientific Reports 12.1 (2022): 1055.

[4] Berrino, Enrico, et al. “Unique patterns of heterogeneous mismatch repair protein expression in colorectal cancer unveil different degrees of tumour mutational burden and distinct tumour microenvironment features.” Modern Pathology 36.2 (2023): 100012.

[5] Garcia, Paulo, et al. “CD8+ T-cell Density Is an Independent Predictor of Survival and Response to Adjuvant Chemotherapy in Stage III Colon Cancer.” Applied Immunohistochemistry & Molecular Morphology 31.2 (2023): 69–76.

[6] Williams, Christopher, et al. “Artificial intelligence-assisted evaluation of tumour infiltrating CD3+ and CD8+ T cells for prognostication and prediction of benefit from adjuvant chemotherapy in early stage colorectal cancer (CRC): A retrospective analysis of the QUASAR trial.” (2023): 204–204.

[7] Bruni, Daniela, Helen K. Angell, and Jérôme Galon. “The immune contexture and Immunoscore in cancer prognosis and therapeutic efficacy.” Nature Reviews Cancer 20.11 (2020): 662–680.

[8] Galon, Jérôme, et al. “Towards the introduction of the ‘Immunoscore’in the classification of malignant tumours.” The Journal of pathology 232.2 (2014): 199–209.

[9] Pagès, Franck, et al. “International validation of the consensus Immunoscore for the classification of colon cancer: a prognostic and accuracy study.” The Lancet 391.10135 (2018): 2128–2139.

[10] Mlecnik, Bernhard, et al. “Integrative analyses of colorectal cancer show immunoscore is a stronger predictor of patient survival than microsatellite instability.” Immunity 44.3 (2016): 698–711.

[11] Li, Chongwu, et al. “The predictive value of inflammatory biomarkers for major pathological response in non-small cell lung cancer patients receiving neoadjuvant chemoimmunotherapy and its association with the immune-related tumour microenvironment: a multi-center study.” Cancer Immunology, Immunotherapy 72.3 (2023): 783–794.

[12] Abdelfatah, Eihab, et al. “Predictive and Prognostic Implications of Circulating CX3CR1+ CD8+ T Cells in Non–Small Cell Lung Cancer Patients Treated with Chemo-Immunotherapy.” Cancer Research Communications 3.3 (2023): 510–520.

[13] Liu, Zhaopei, et al. “Intratumoural TIGIT+ CD8+ T-cell infiltration determines poor prognosis and immune evasion in patients with muscle-invasive bladder cancer.” Journal for immunotherapy of cancer 8.2 (2020).

[14] Giraldo, Nicolas A., et al. “Orchestration and prognostic significance of immune checkpoints in the microenvironment of primary and metastatic renal cell cancer.” Clinical Cancer Research 21.13 (2015): 3031–3040.

[15] Choi, Paul J., and Timothy J. Mitchison. “Imaging burst kinetics and spatial coordination during serial killing by single natural killer cells.” Proceedings of the National Academy of Sciences 110.16 (2013): 6488–6493.

[16] Manaster, Yoav, et al. “Reduced CTL motility and activity in avascular tumour areas.” Cancer Immunology, Immunotherapy 68 (2019): 1287–1301.

[17] Weigelin, Bettina, et al. “Cytotoxic T cells are able to efficiently eliminate cancer cells by additive cytotoxicity.” Nature communications 12.1 (2021): 5217.

[18] Summers, Huw D., John W. Wills, and Paul Rees. “Spatial statistics is a comprehensive tool for quantifying cell neighbor relationships and biological processes via tissue image analysis.” Cell Reports Methods 2.11 (2022).

[19] Gerdes, Michael J., et al. “Highly multiplexed single-cell analysis of formalin-fixed, paraffin-embedded cancer tissue.” Proceedings of the National Academy of Sciences 110.29 (2013): 11982–11987.

[20] Berens, Michael E., et al. “Multiscale, multimodal analysis of tumour heterogeneity in IDH1 mutant vs wild-type diffuse gliomas.” PLoS One 14.12 (2019): e0219724.

[21] Stachtea, Xanthi, et al. “Stratification of chemotherapy-treated stage III colorectal cancer patients using multiplexed imaging and single-cell analysis of T-cell populations.” Modern Pathology 35.4 (2022): 564–576.

[22] Kumar, G., Bangi, M., An Extension to Winding Number and Point-in-Polygon Algorithm. IFAC-PapersOnLine 51, 1 (2018).

[23] Wang, Xiao Qian, et al. “Spatial predictors of immunotherapy response in triple-negative breast cancer.” Nature 621.7980 (2023): 868–876.

[24] Failmezger, H. et al. “Topological tumor graphs: a graph-based spatial model to infer stromal recruitment for immunosuppression in melanoma histology”. Cancer Research 80.5, (2020) : 1199–1209.

[25] Najem, H. et al. “Central nervous system immune interactome is a function of cancer lineage, tumor microenvironment, and STAT3 expression.” JCI insight, 7.9 (2022).

[26] Lindner, A. U. et al. “An atlas of inter-and intra-tumor heterogeneity of apoptosis competency in colorectal cancer tissue at single-cell resolution.” Cell Death & Differentiation, 29.4 (2022): 806–817.

[27] Fluge, Ø., et al. “Expression of EZH2 and Ki-67 in colorectal cancer and associations with treatment response and prognosis.” British journal of cancer 101.8 (2009): 1282–1289.

[28] Kim, Tae Hyun, et al. “GLUT-1 may predict metastases and death in patients with locally advanced rectal cancer.” Frontiers in Oncology 13 (2023): 1094480.

