## Supplementary figures and images for "Spatial Effects of Infiltrating T cells on Neighbouring Cancer Cells and Prognosis in Stage III CRC patients"

### Suppl-fig-1

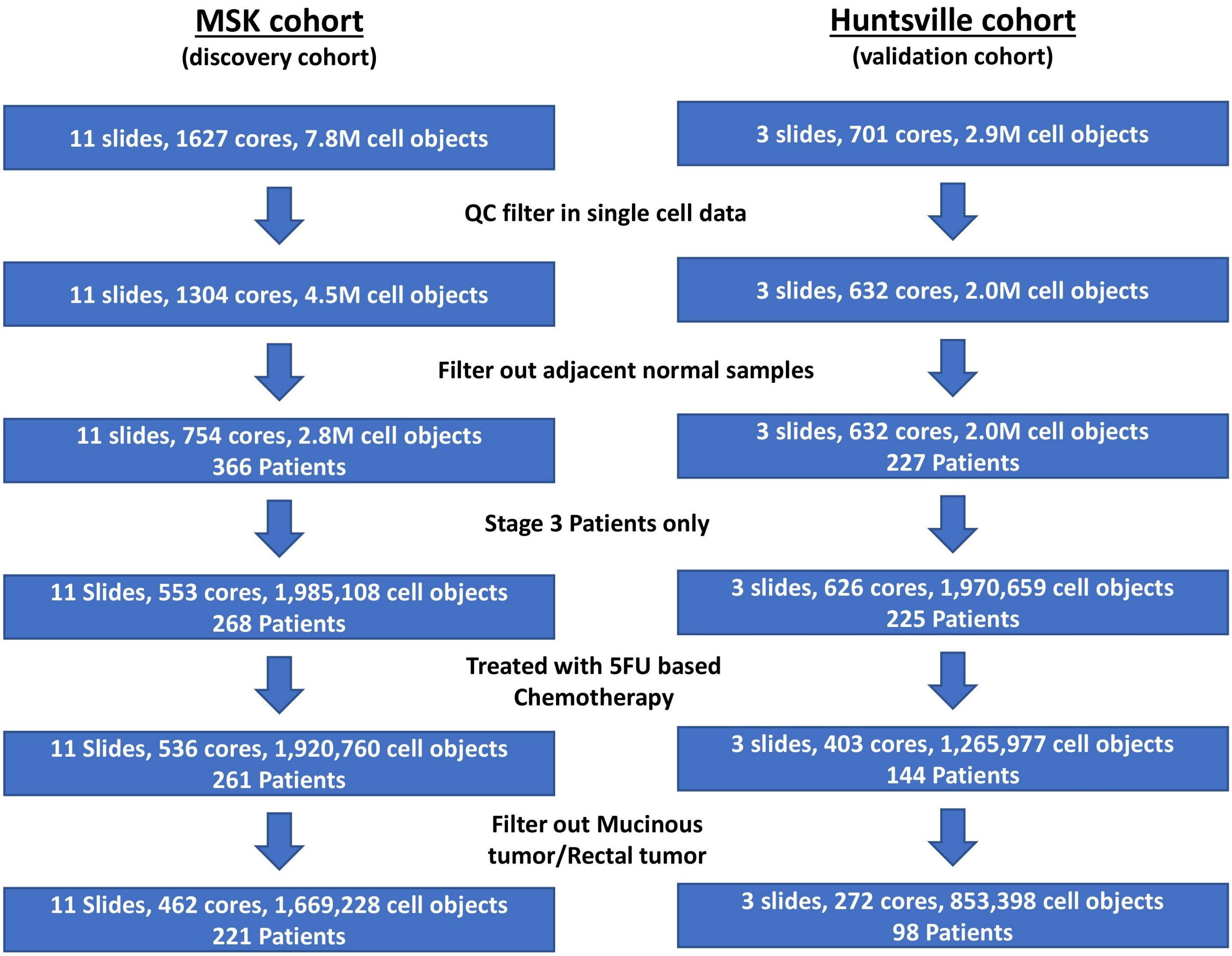

### Suppl-fig-3

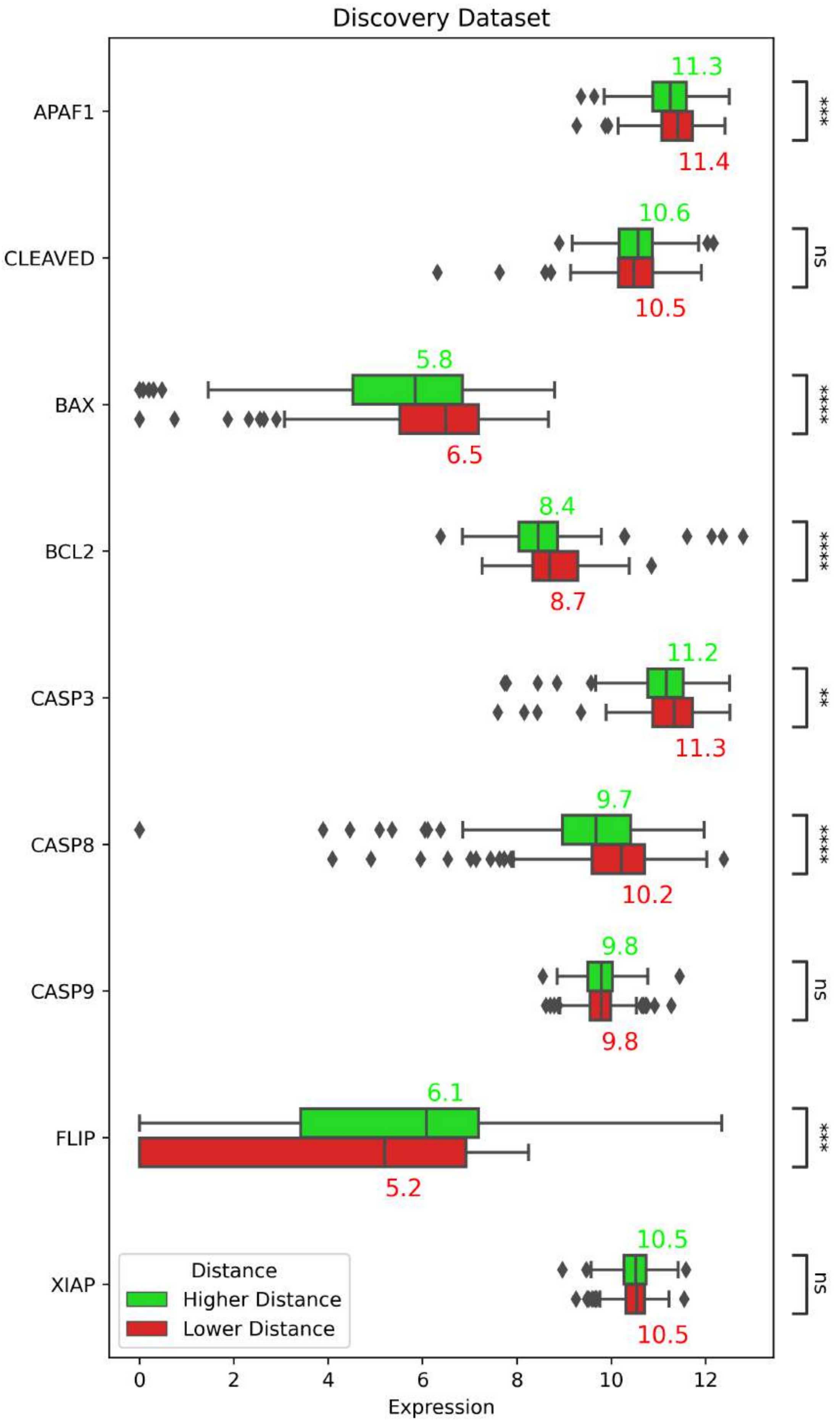

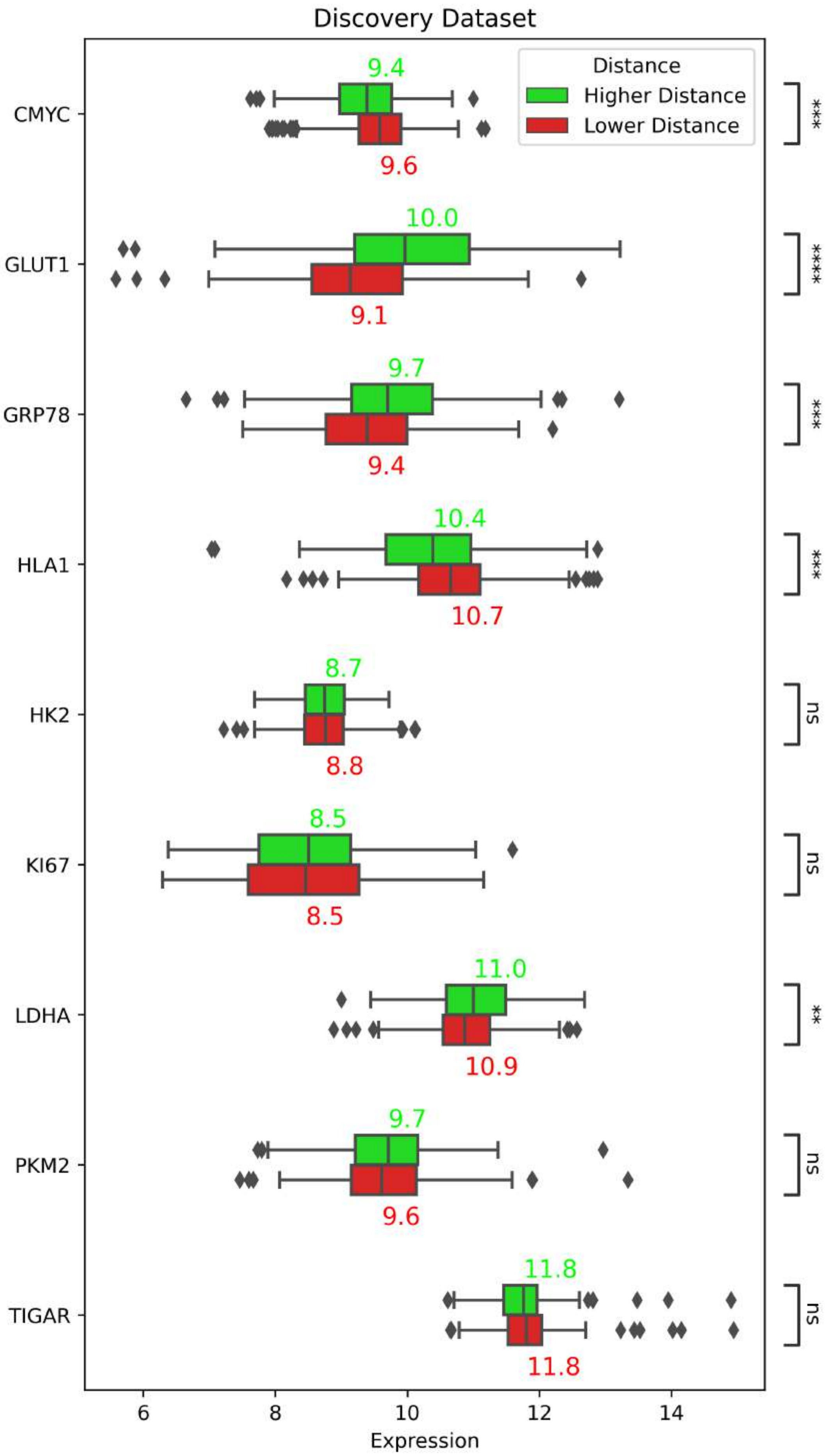

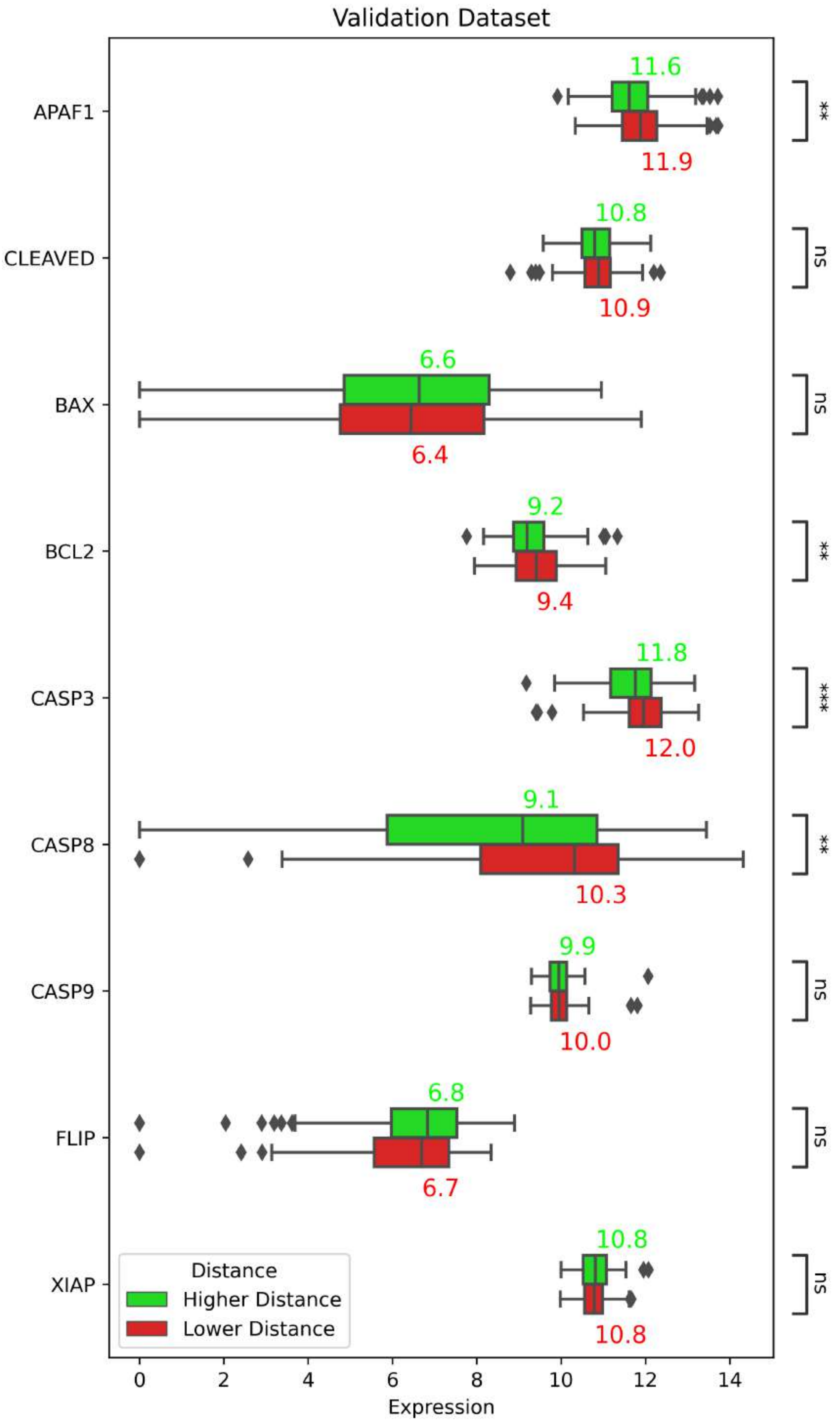

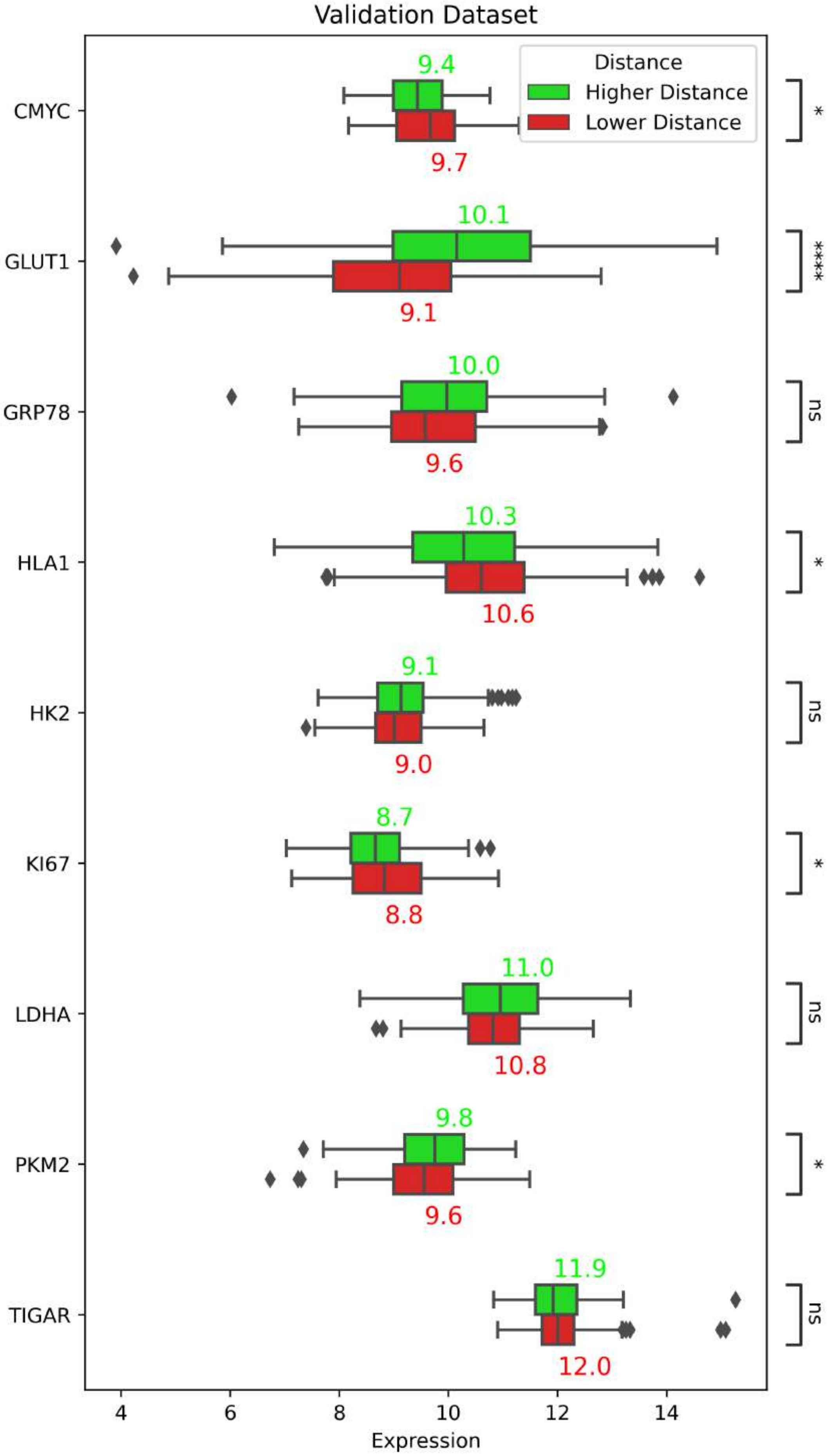

### Suppl-fig-4

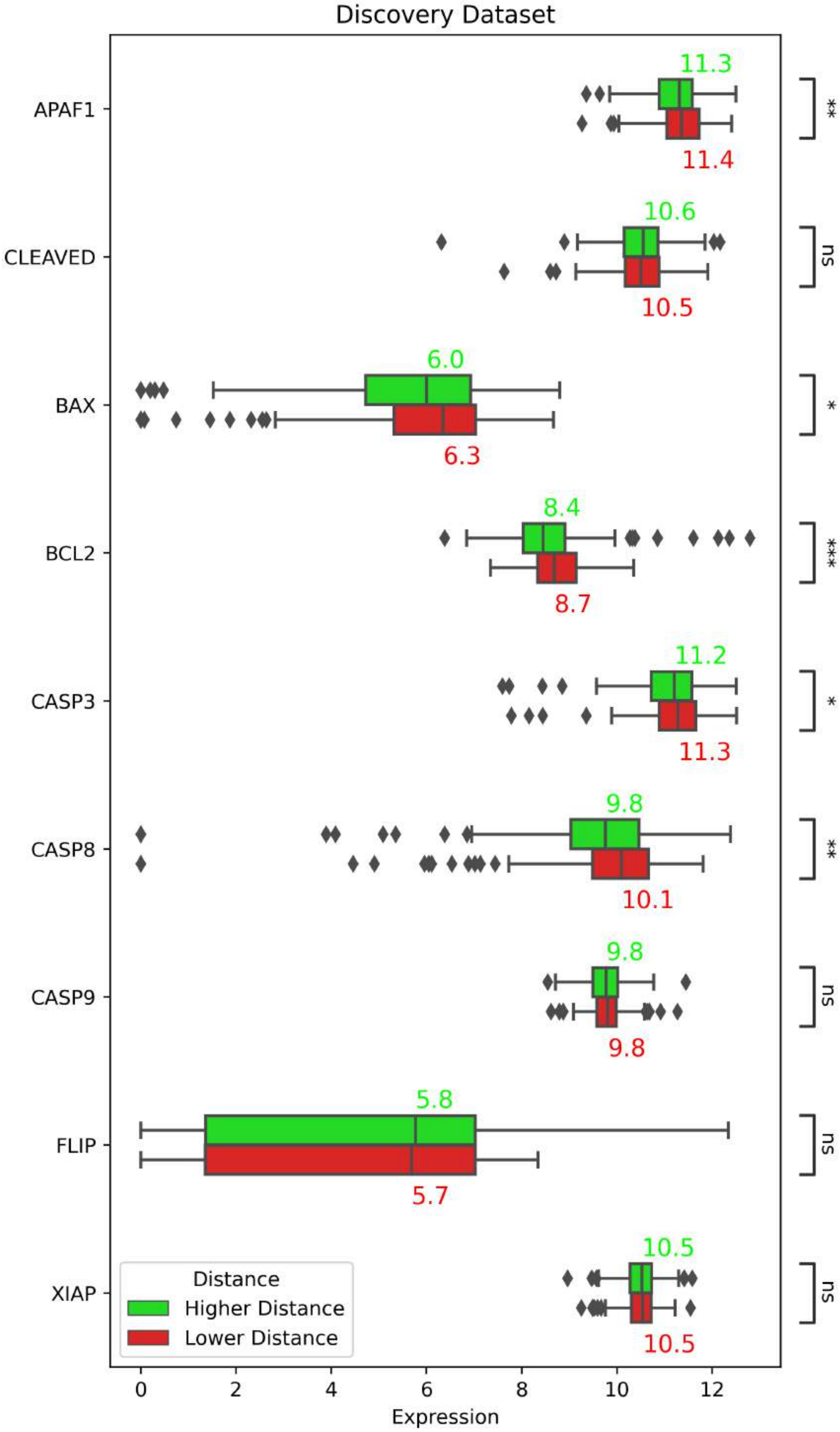

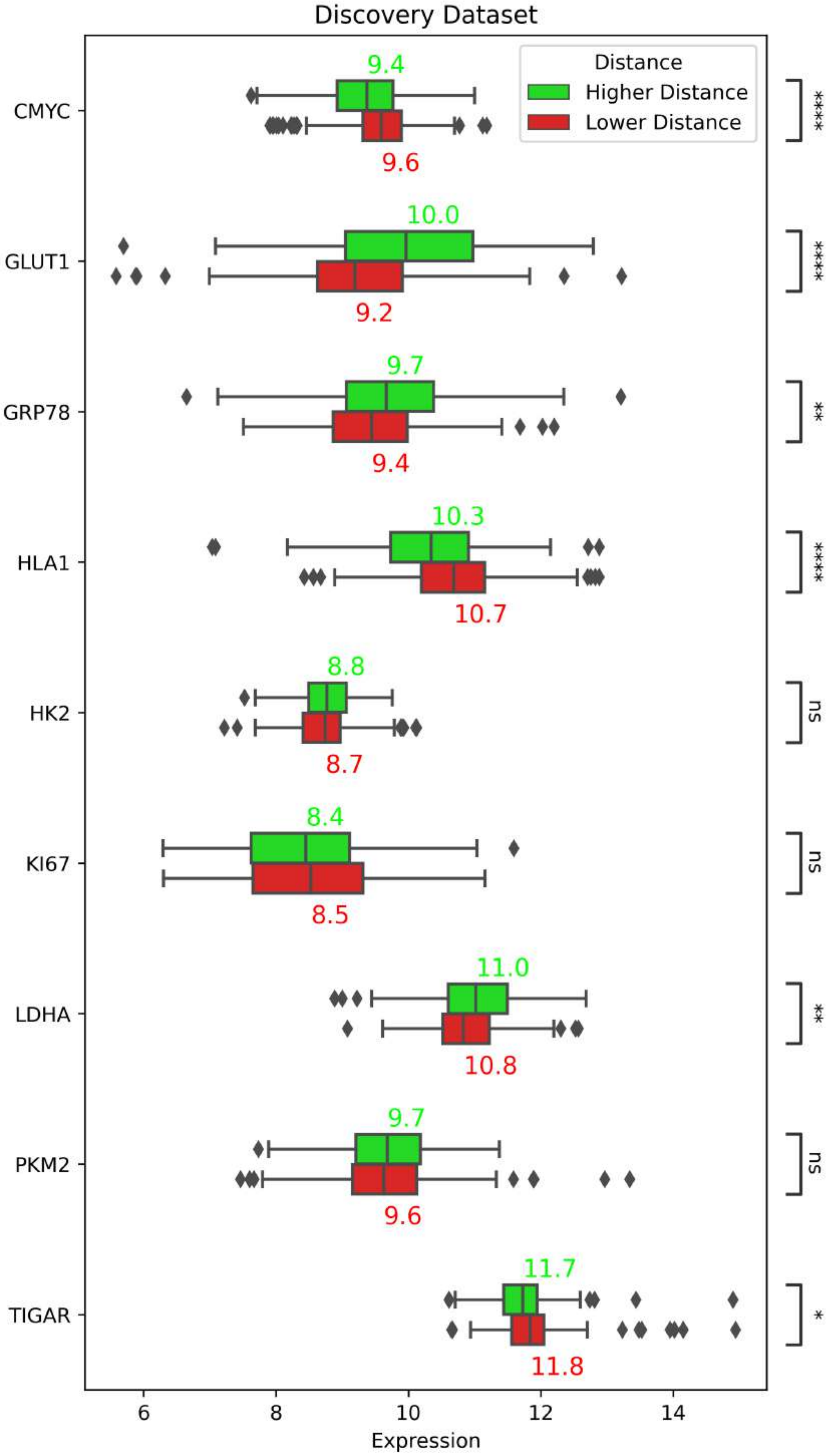

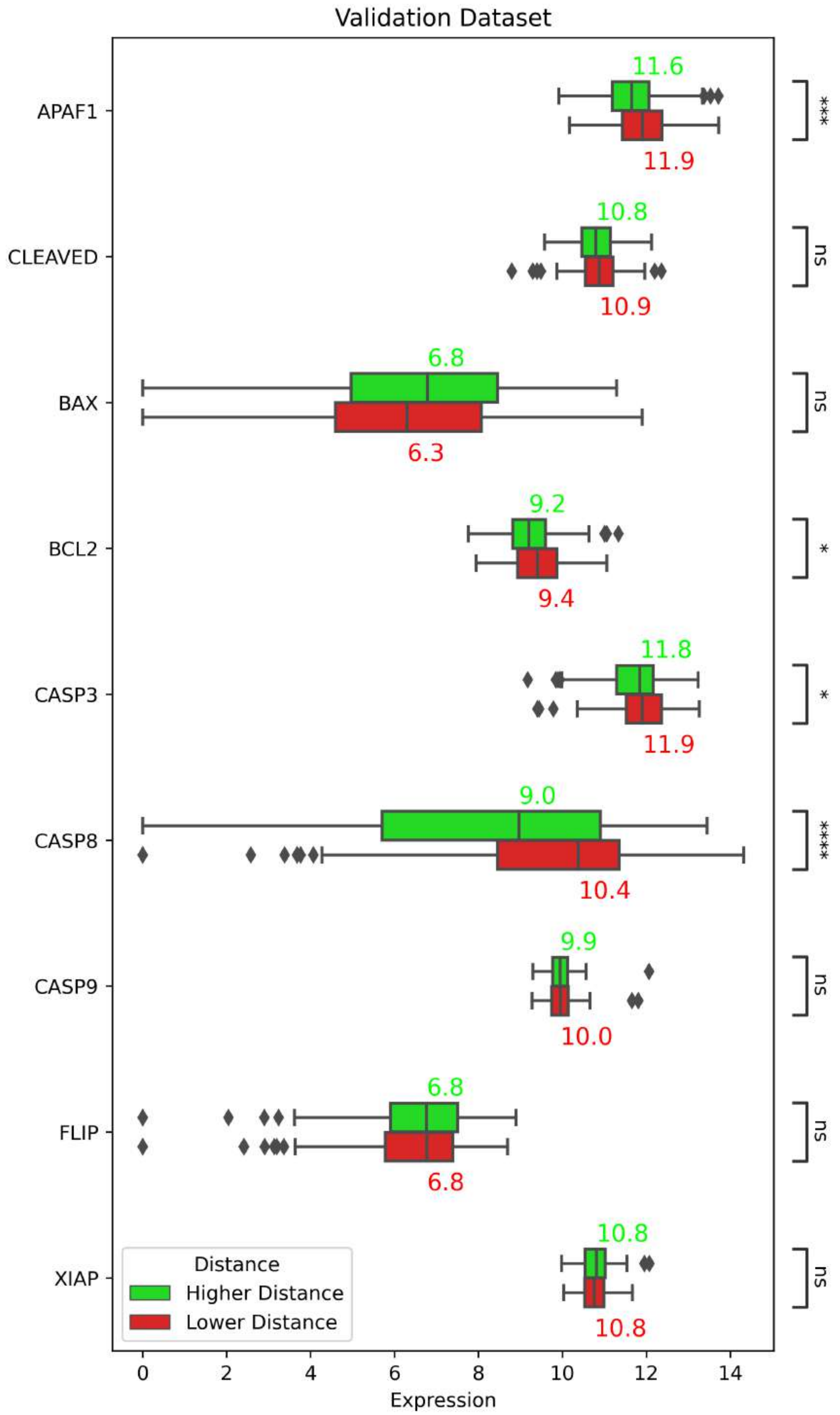

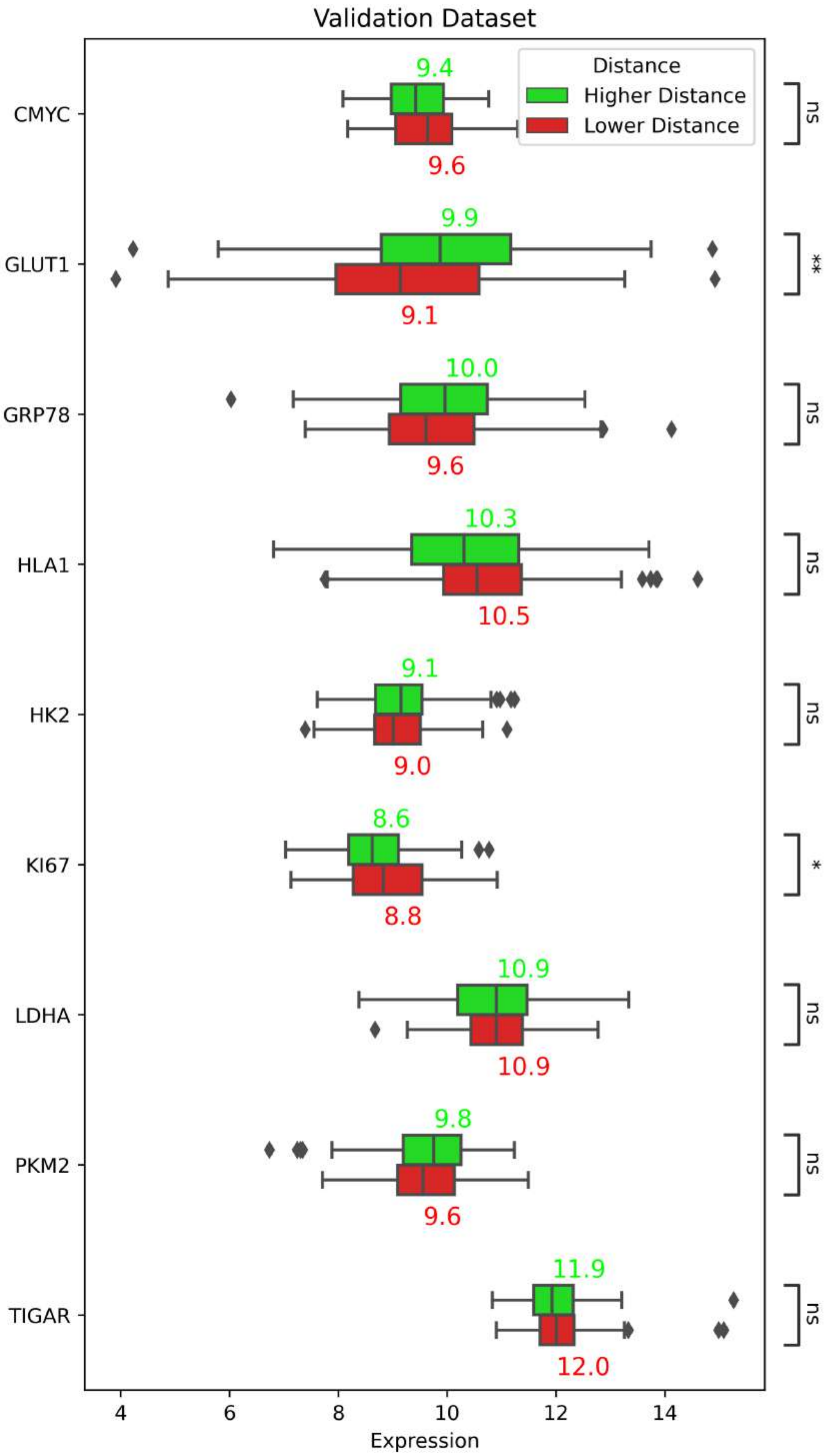

### Suppl-fig-5

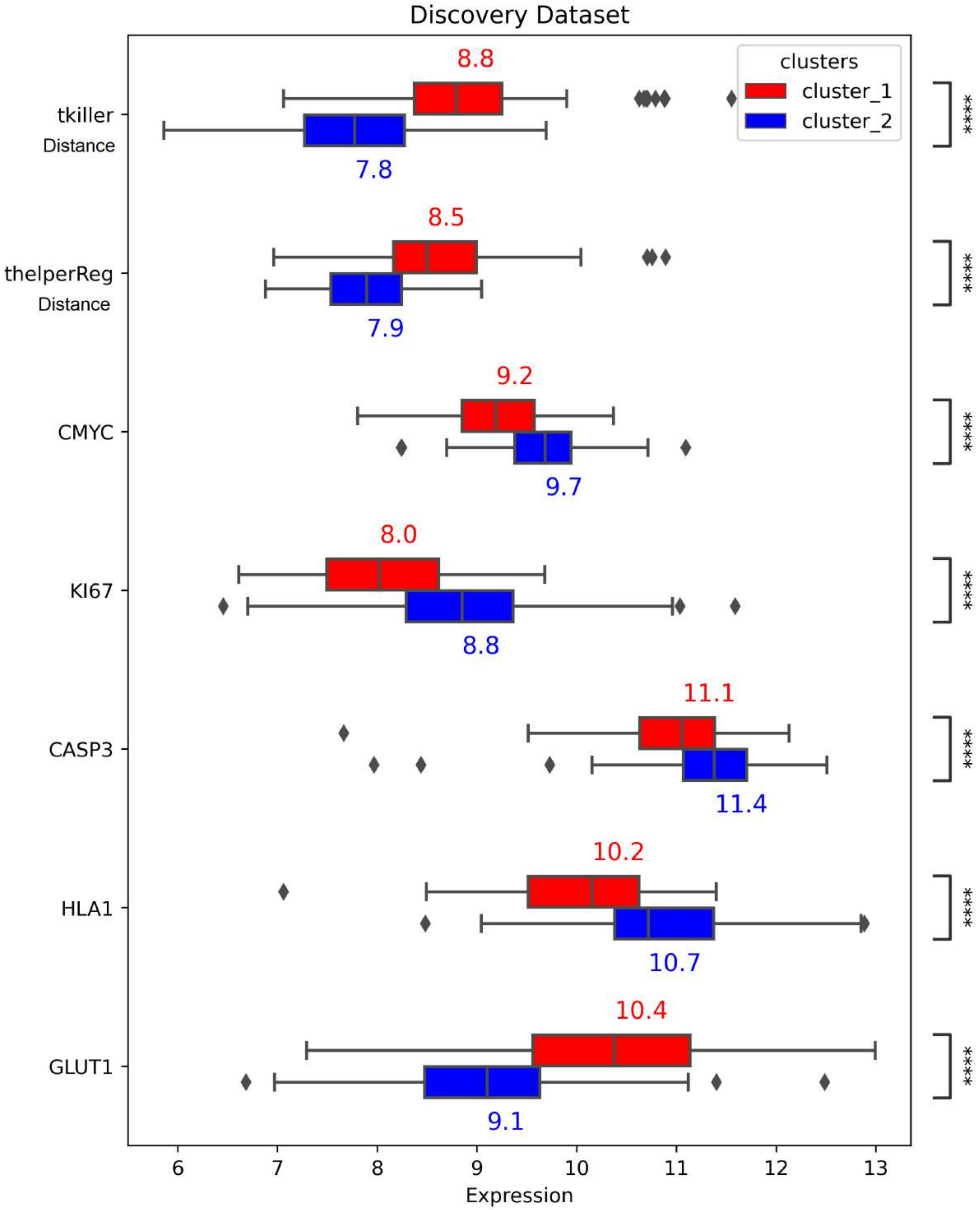

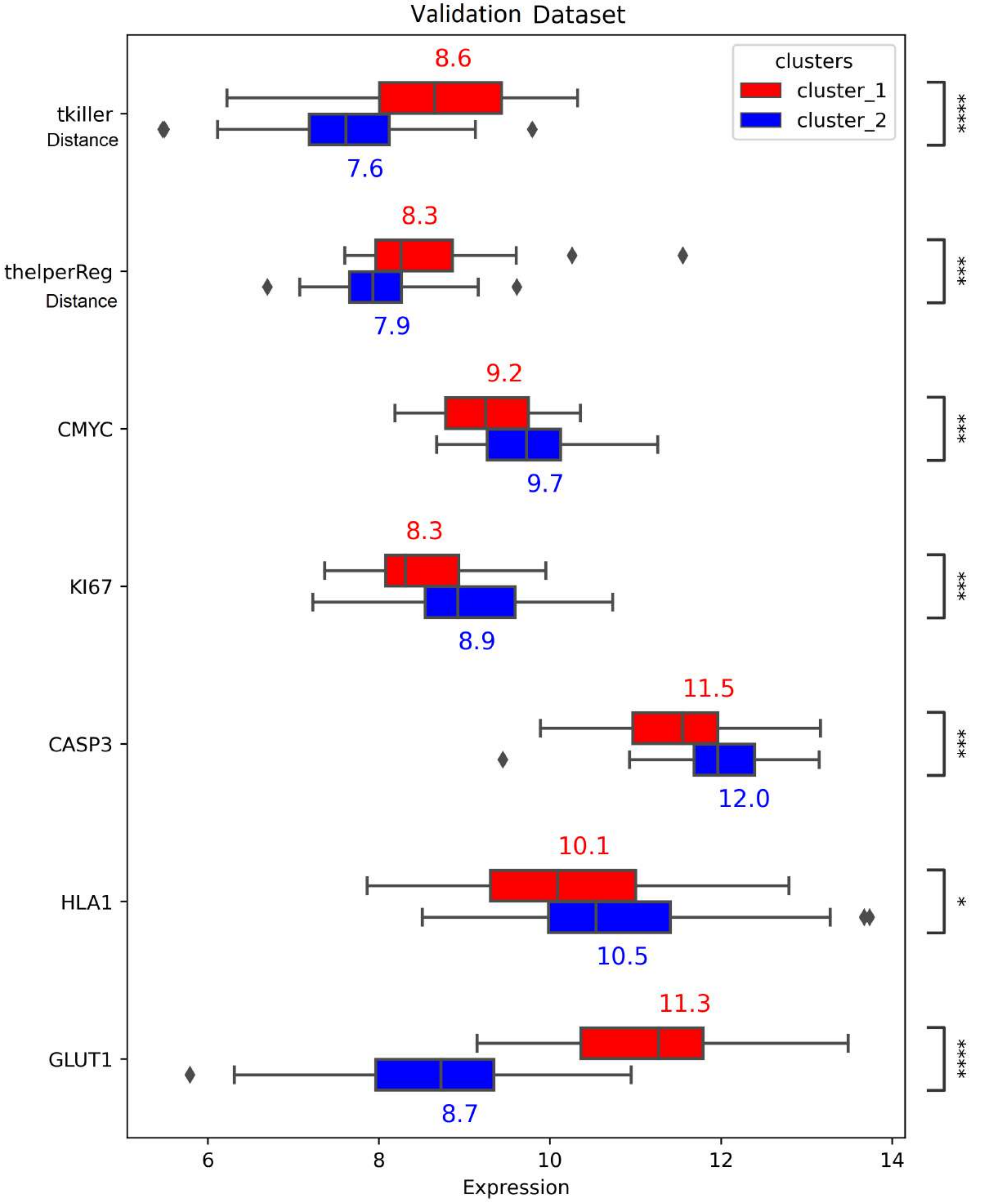

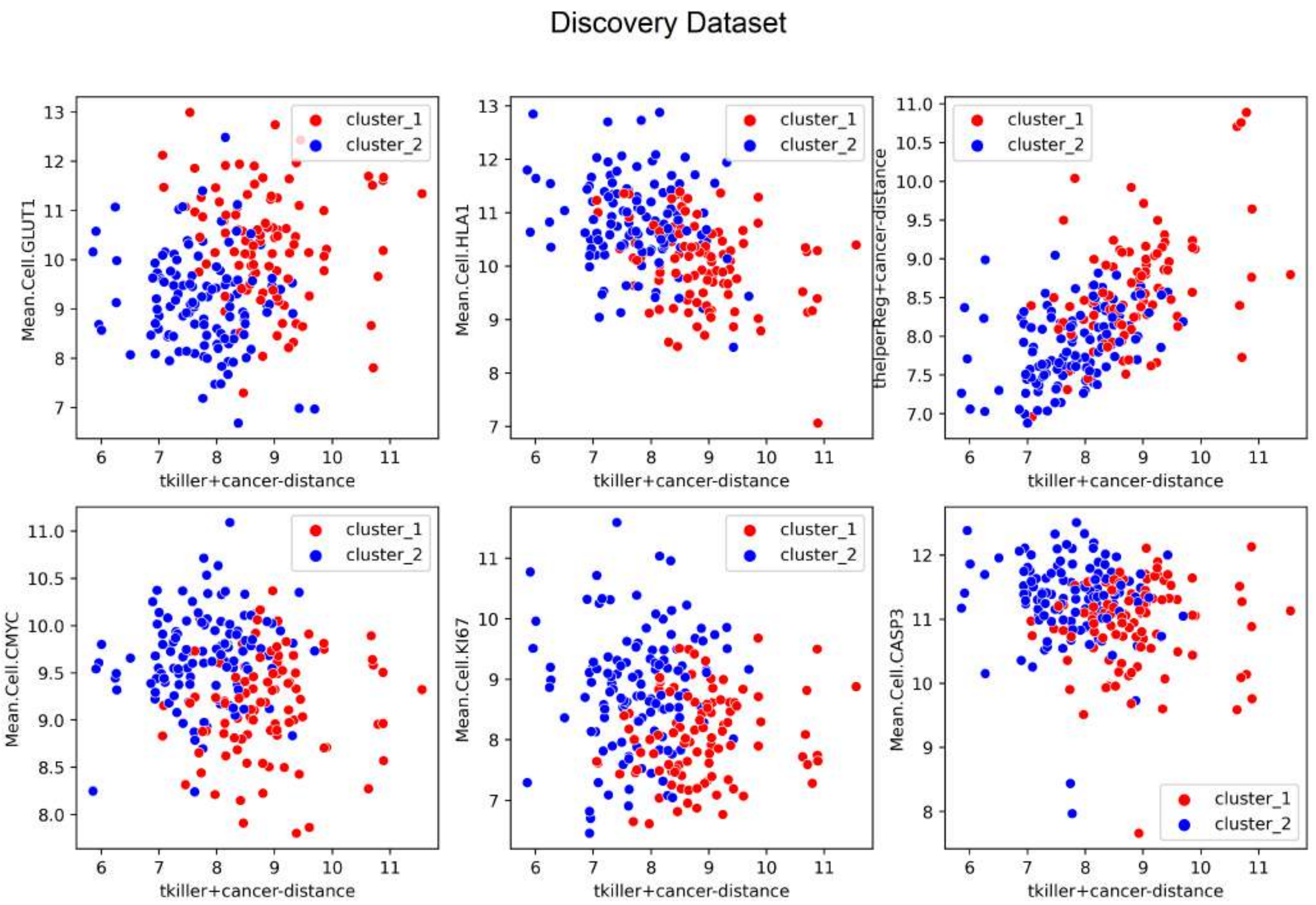

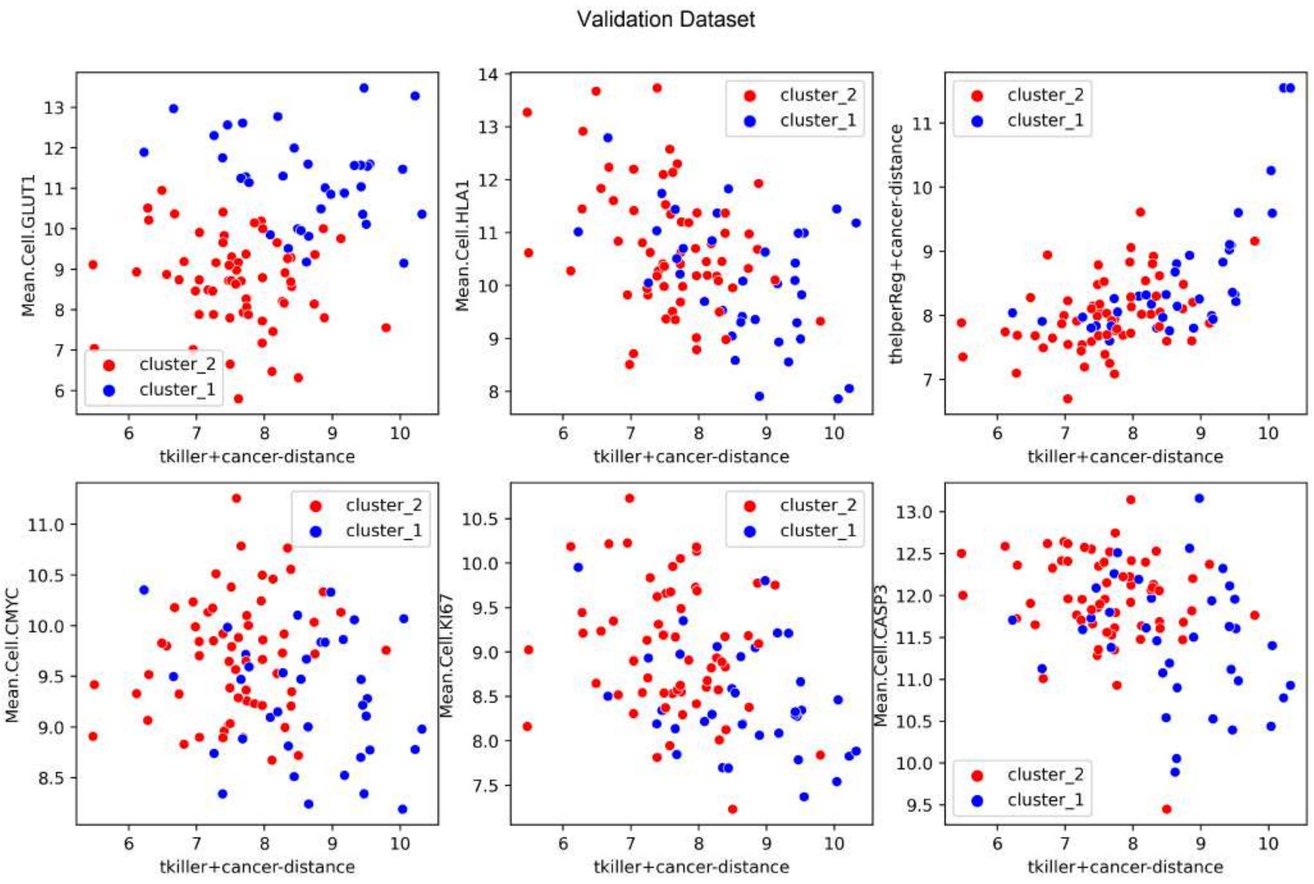

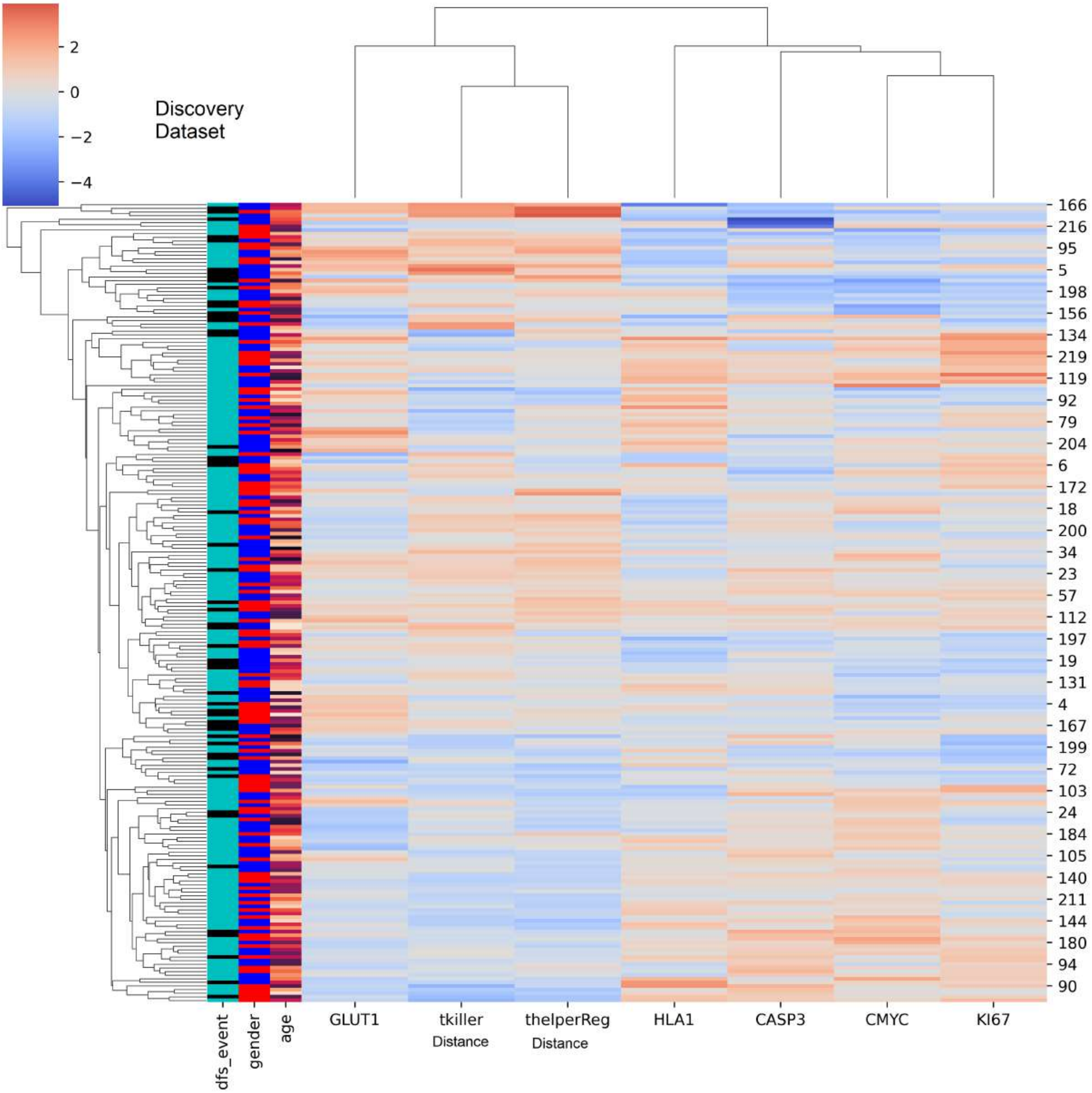

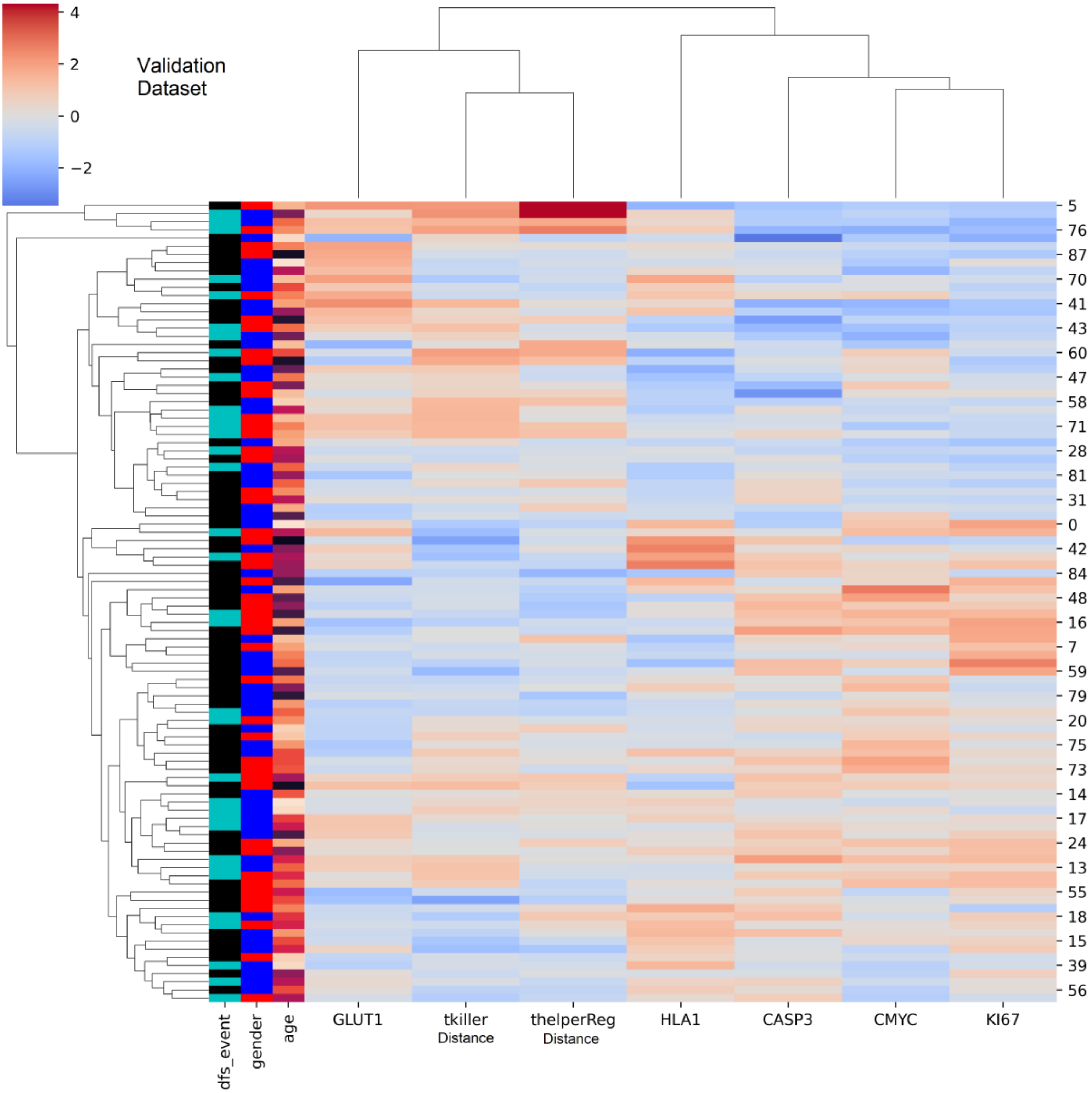
