## Supplementary material for "Spatial Effects of Infiltrating T cells on Neighbouring Cancer Cells and Prognosis in Stage III CRC patients": Suppl-fig-2

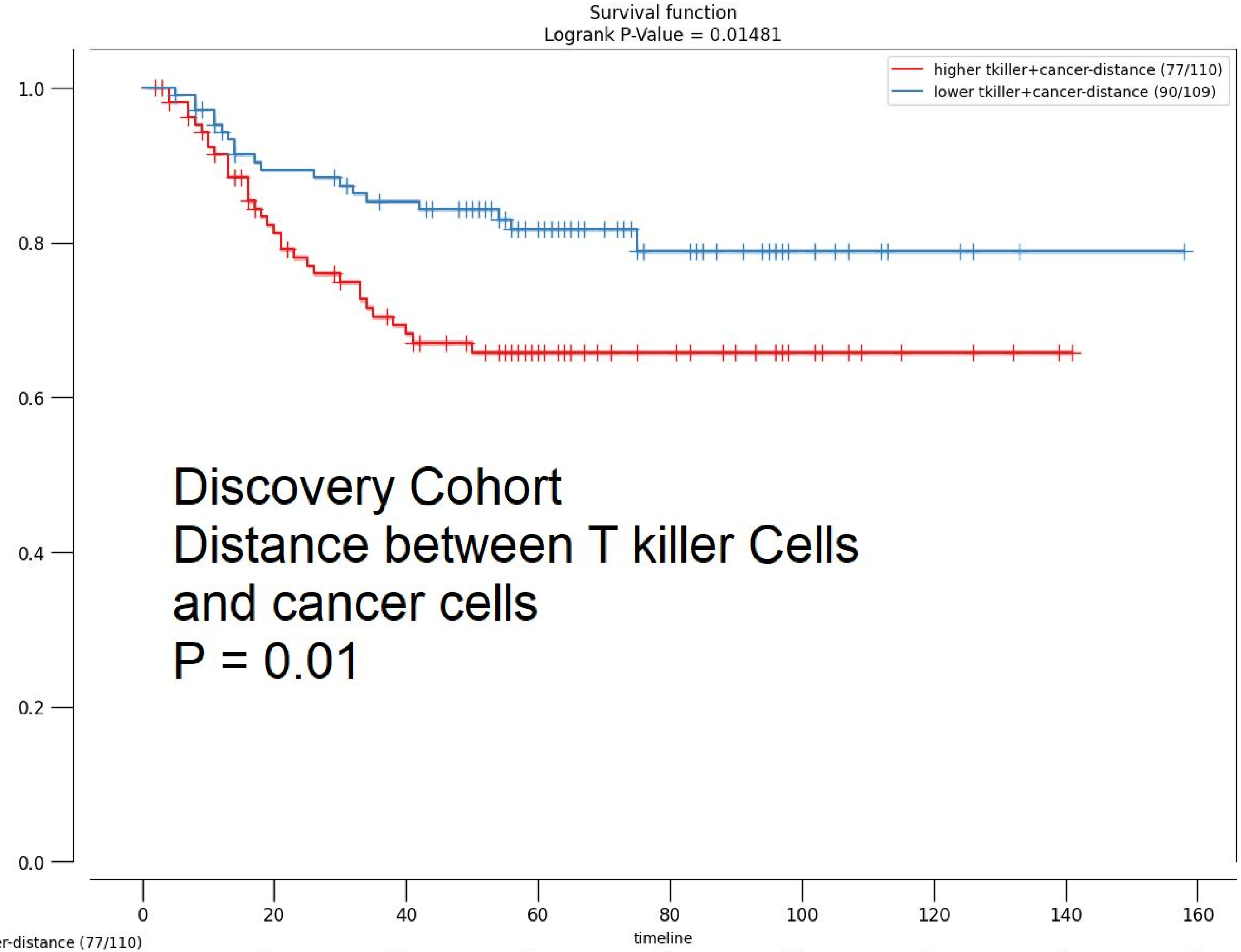


| | 0 | 20 | 40 | 60 | 80 | 100 | 120 | 140 | 160 |
|---|---|---|---|---|---|---|---|---|---|
| higher tkiller+cancer-distance (77/110) | | | | | | | | | |
| At risk | 110 | 78 | 60 | 34 | 19 | 10 | 4 | 1 | 0 |
| Censored | 0 | 13 | 19 | 43 | 58 | 67 | 73 | 76 | 77 |
| Events | 0 | 19 | 31 | 33 | 33 | 33 | 33 | 33 | 33 |
| lower tkiller+cancer-distance (90/109) | | | | | | | | | |
| At risk | 109 | 89 | 82 | 52 | 25 | 9 | 4 | 1 | 0 |
| Censored | 0 | 9 | 12 | 39 | 65 | 81 | 86 | 89 | 90 |
| Events | 0 | 11 | 15 | 18 | 19 | 19 | 19 | 19 | 19 |

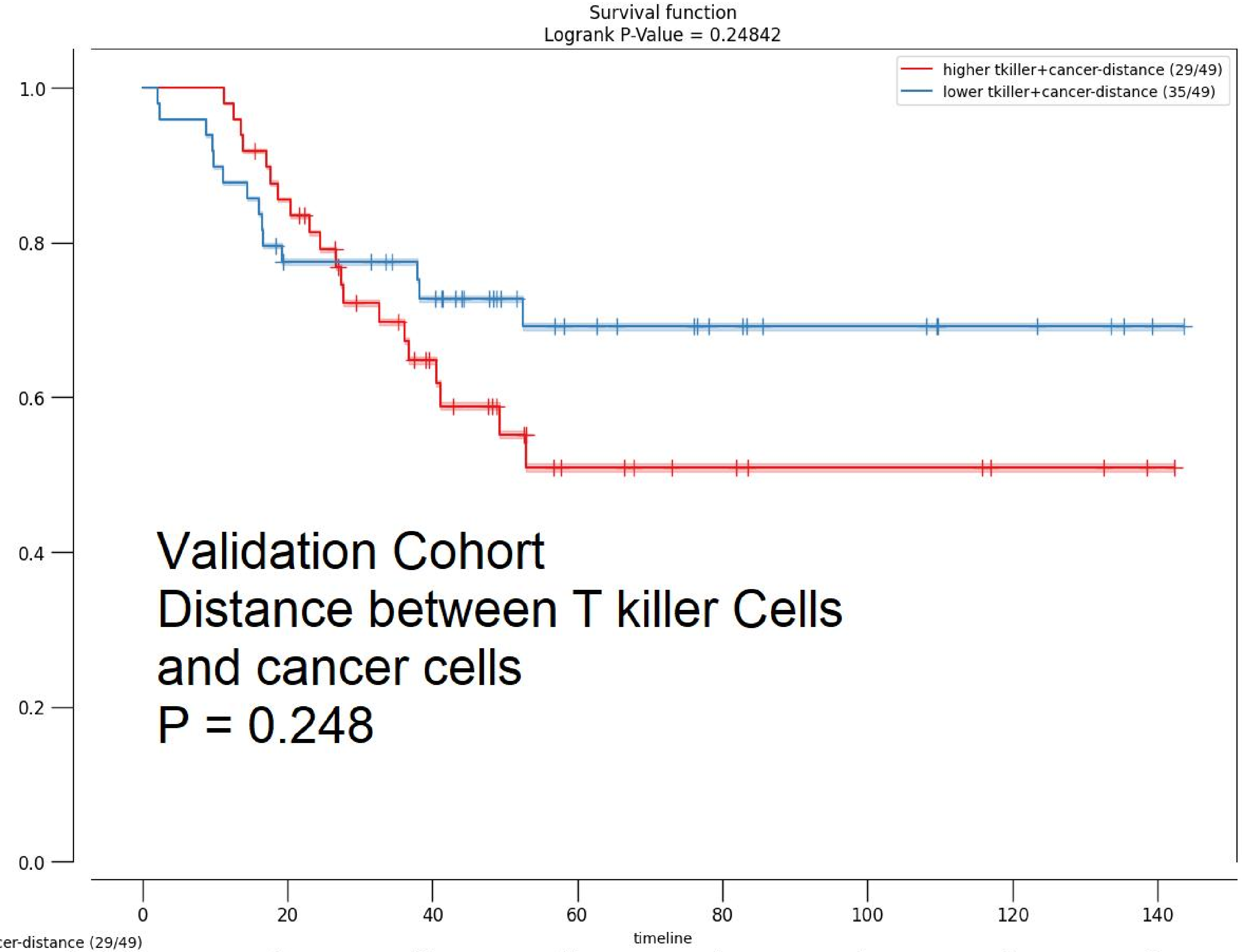


| | 0 | 20 | 40 | 60 | 80 | 100 | 120 | 140 |
|---|---|---|---|---|---|---|---|---|
| higher tkiller+cancer-distance (29/49) | | | | | | | | |
| At risk | 49 | 41 | 22 | 10 | 7 | 5 | 3 | 1 |
| Censored | 0 | 1 | 11 | 19 | 22 | 24 | 26 | 28 |
| Events | 0 | 7 | 16 | 20 | 20 | 20 | 20 | 20 |
| lower tkiller+cancer-distance (35/49) | | | | | | | | |
| At risk | 49 | 36 | 31 | 17 | 11 | 8 | 5 | 1 |
| Censored | 0 | 2 | 5 | 18 | 24 | 27 | 30 | 34 |
| Events | 0 | 11 | 13 | 14 | 14 | 14 | 14 | 14 |

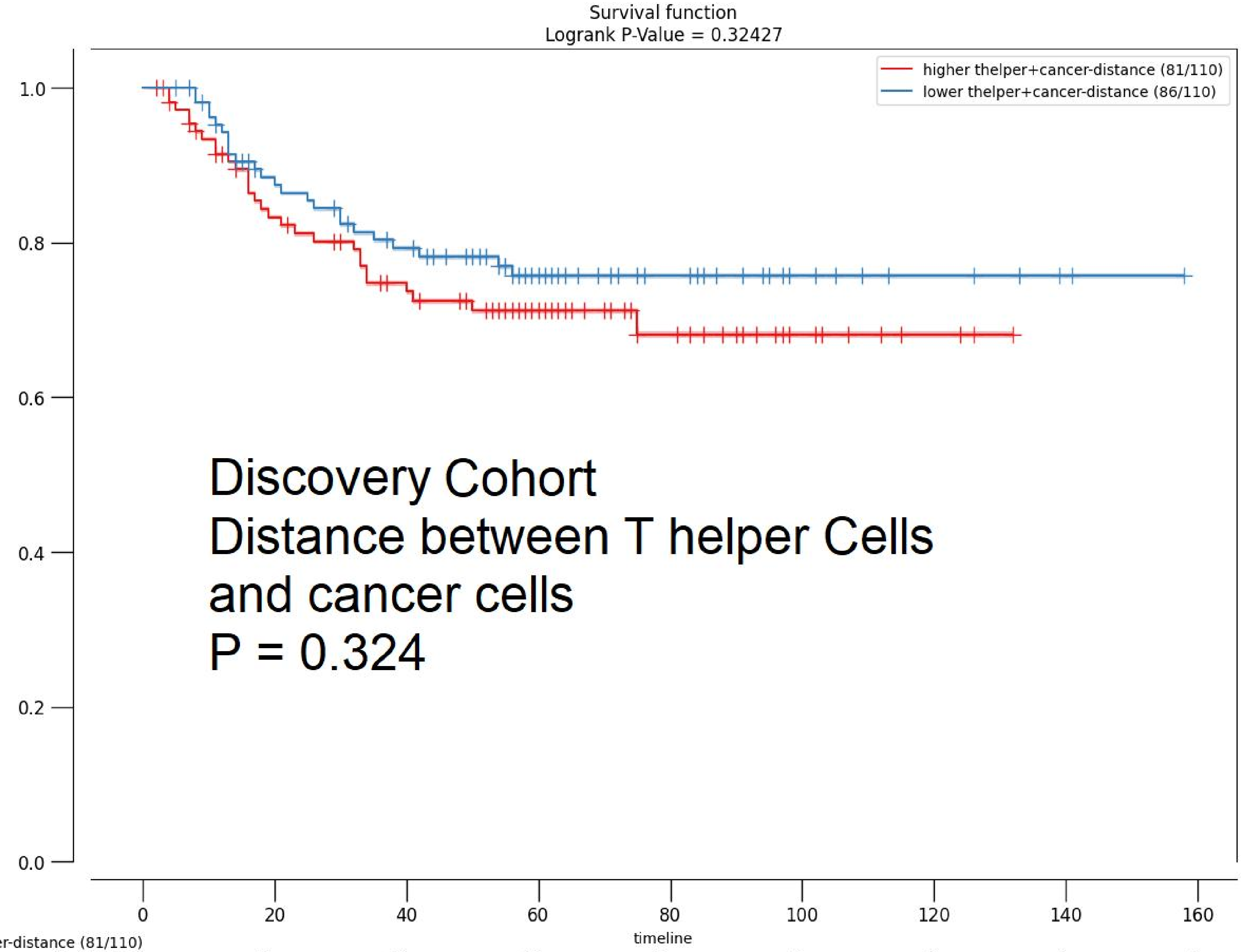


| | 0 | 20 | 40 | 60 | 80 | 100 | 120 | 140 | 160 |
|---|---|---|---|---|---|---|---|---|---|
| **higher thelper+cancer-distance (81/110)** | | | | | | | | | |
| At risk | 110 | 81 | 66 | 42 | 21 | 9 | 3 | 0 | 0 |
| Censored | 0 | 12 | 18 | 40 | 60 | 72 | 78 | 81 | 81 |
| Events | 0 | 17 | 26 | 28 | 29 | 29 | 29 | 29 | 29 |
| **lower thelper+cancer-distance (86/110)** | | | | | | | | | |
| At risk | 110 | 87 | 76 | 44 | 23 | 10 | 5 | 2 | 0 |
| Censored | 0 | 10 | 13 | 42 | 63 | 76 | 81 | 84 | 86 |
| Events | 0 | 13 | 21 | 24 | 24 | 24 | 24 | 24 | 24 |

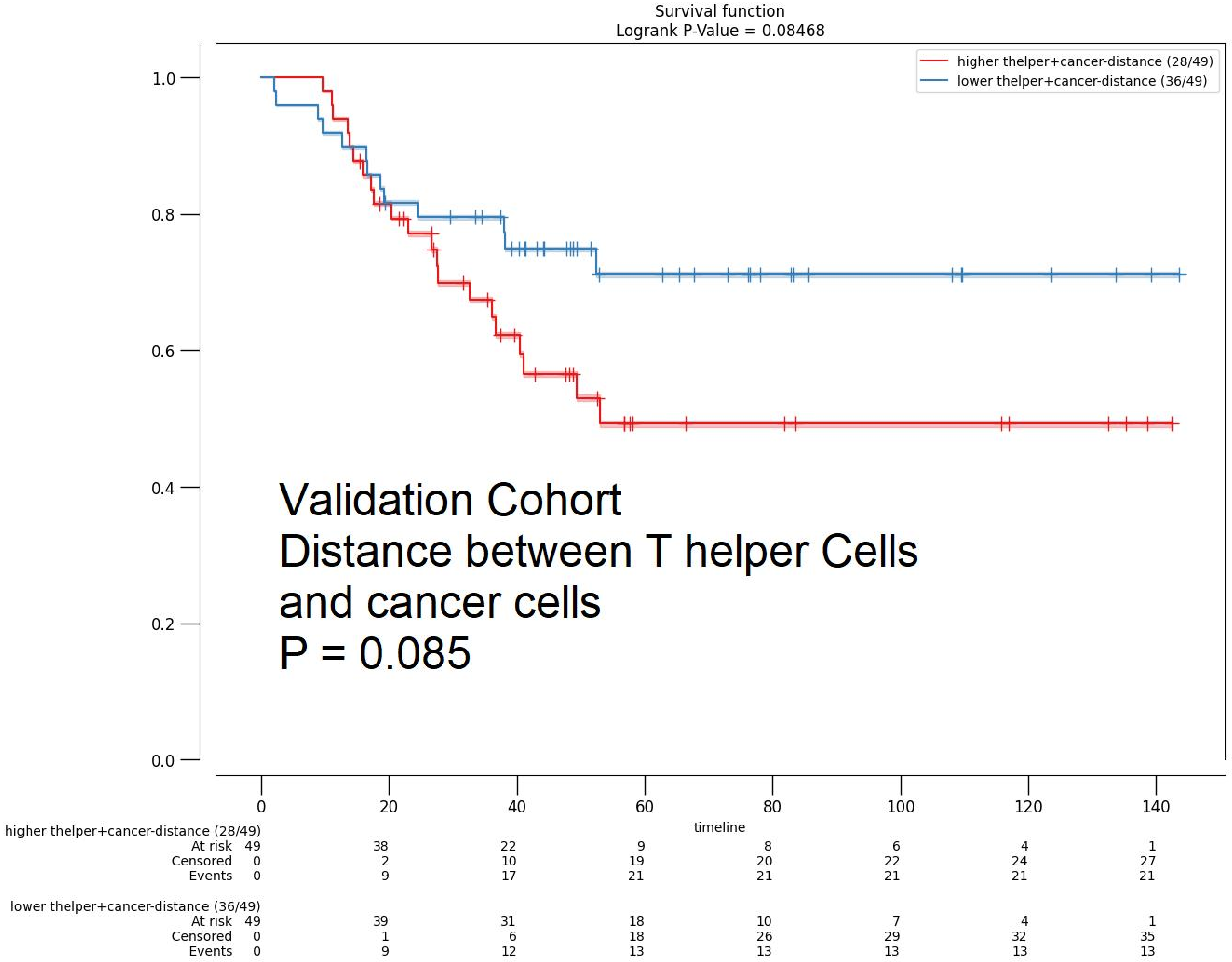
Survival function
Logrank P-Value = 0.08468
higher thelper+cancer-distance (28/49)
lower thelper+cancer-distance (36/49)
1.0
0.8
0.6
0.4
0.2
0.0
Validation Cohort
Distance between T helper Cells
and cancer cells
P = 0.085
0
20
40
60
80
100
120
140
timeline
higher thelper+cancer-distance (28/49)
At risk 49 38 22 9 8 6 4 1
Censored 0 2 10 19 20 22 24 27
Events 0 9 17 21 21 21 21 21
lower thelper+cancer-distance (36/49)
At risk 49 39 31 18 10 7 4 1
Censored 0 1 6 18 26 29 32 35
Events 0 9 12 13 13 13 13 13

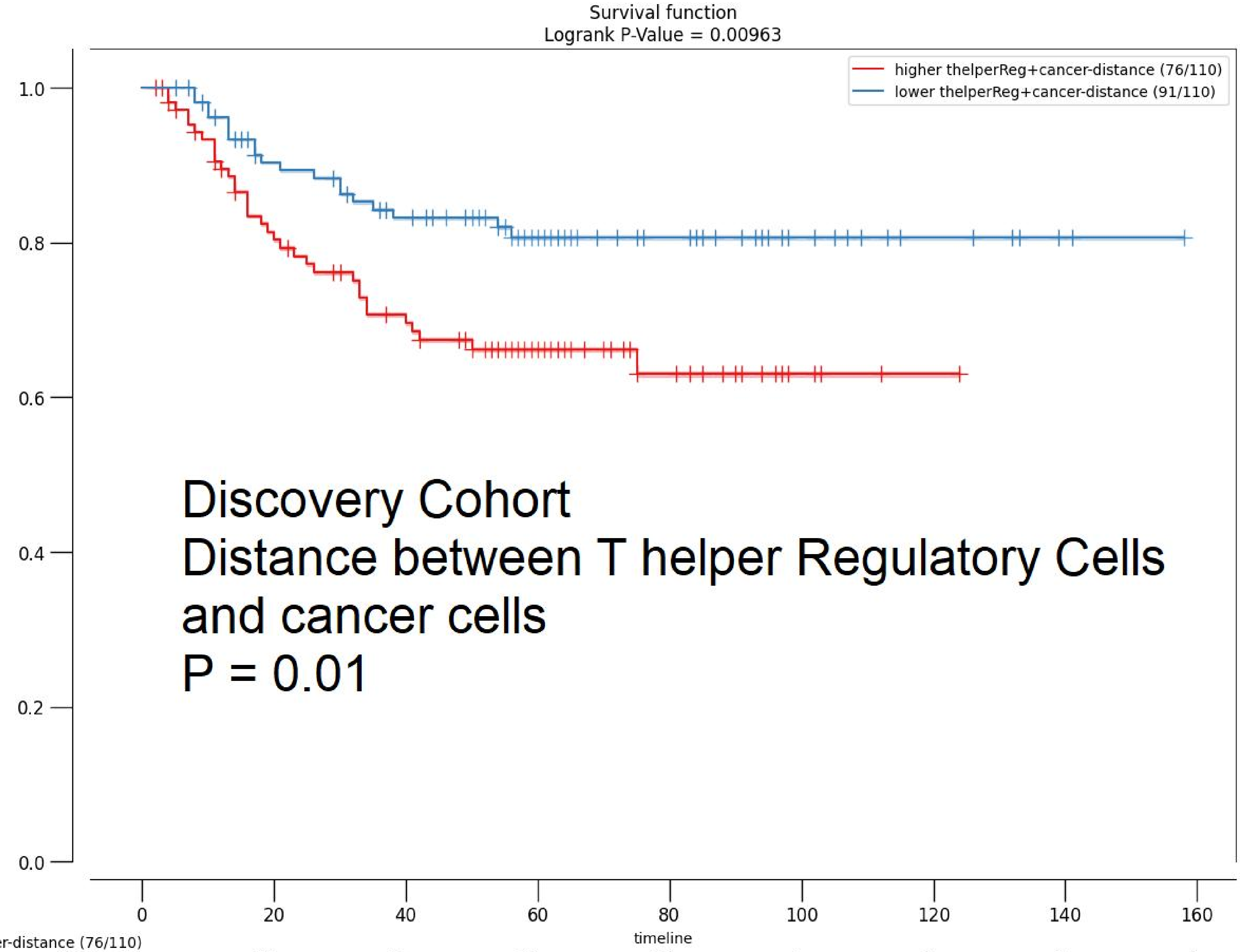


| | 0 | 20 | 40 | 60 | 80 | 100 | 120 | 140 | 160 |
|---|---|---|---|---|---|---|---|---|---|
| **higher thelperReg+cancer-distance (76/110)** | | | | | | | | | |
| At risk | 110 | 78 | 63 | 39 | 18 | 4 | 1 | 0 | 0 |
| Censored | 0 | 12 | 17 | 38 | 58 | 72 | 75 | 76 | 76 |
| Events | 0 | 20 | 30 | 33 | 34 | 34 | 34 | 34 | 34 |
| **lower thelperReg+cancer-distance (91/110)** | | | | | | | | | |
| At risk | 110 | 90 | 79 | 47 | 26 | 15 | 7 | 2 | 0 |
| Censored | 0 | 10 | 14 | 44 | 65 | 76 | 84 | 89 | 91 |
| Events | 0 | 10 | 17 | 19 | 19 | 19 | 19 | 19 | 19 |

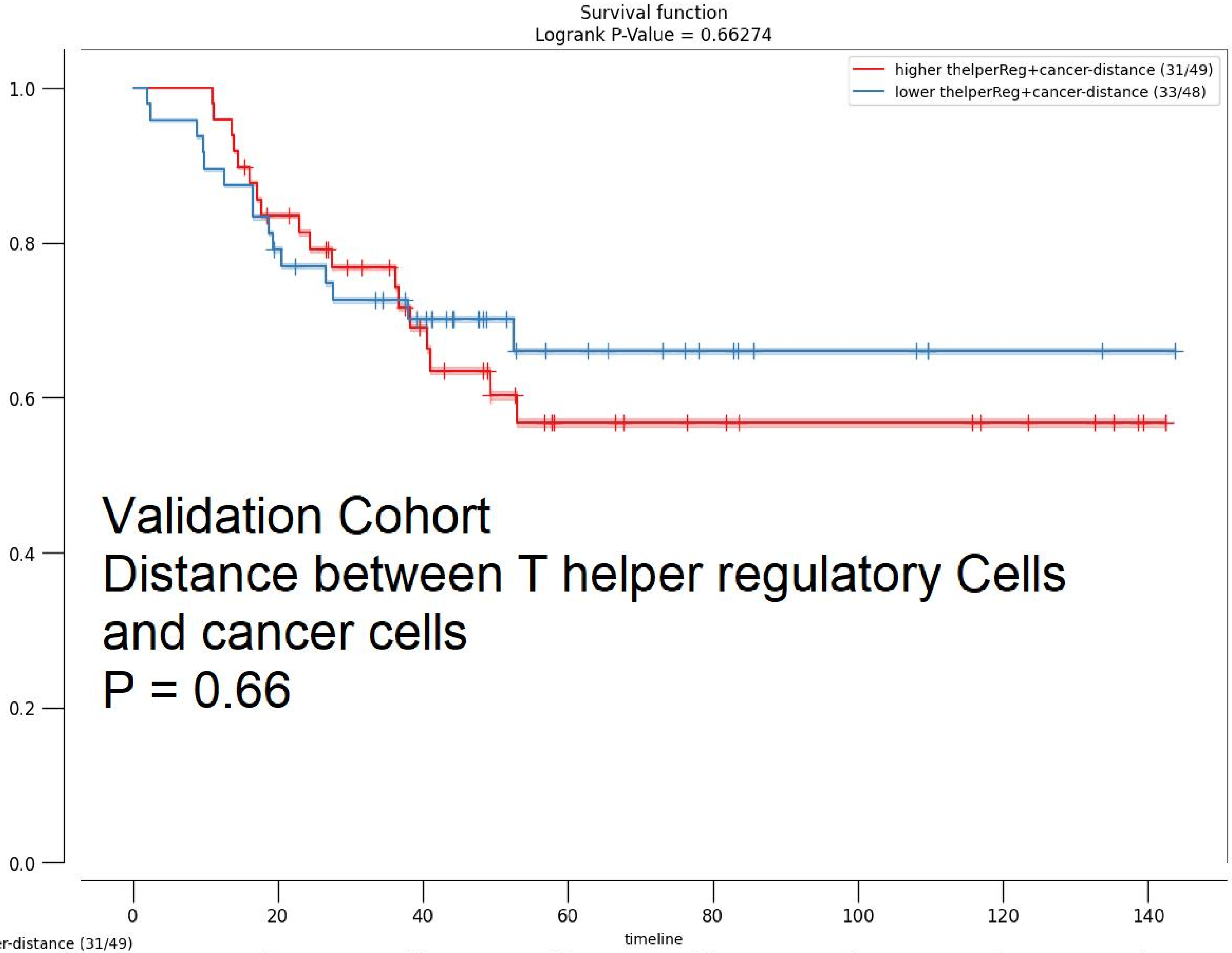


| | 0 | 20 | 40 | 60 | 80 | 100 | 120 | 140 |
|---|---|---|---|---|---|---|---|---|
| higher thelperReg+cancer-distance (31/49) | | | | | | | | |
| At risk | 49 | 39 | 25 | 13 | 10 | 8 | 6 | 1 |
| Censored | 0 | 2 | 10 | 18 | 21 | 23 | 25 | 30 |
| Events | 0 | 8 | 14 | 18 | 18 | 18 | 18 | 18 |
| lower thelperReg+cancer-distance (33/48) | | | | | | | | |
| At risk | 48 | 37 | 28 | 14 | 8 | 5 | 2 | 1 |
| Censored | 0 | 1 | 6 | 19 | 25 | 28 | 31 | 32 |
| Events | 0 | 10 | 14 | 15 | 15 | 15 | 15 | 15 |

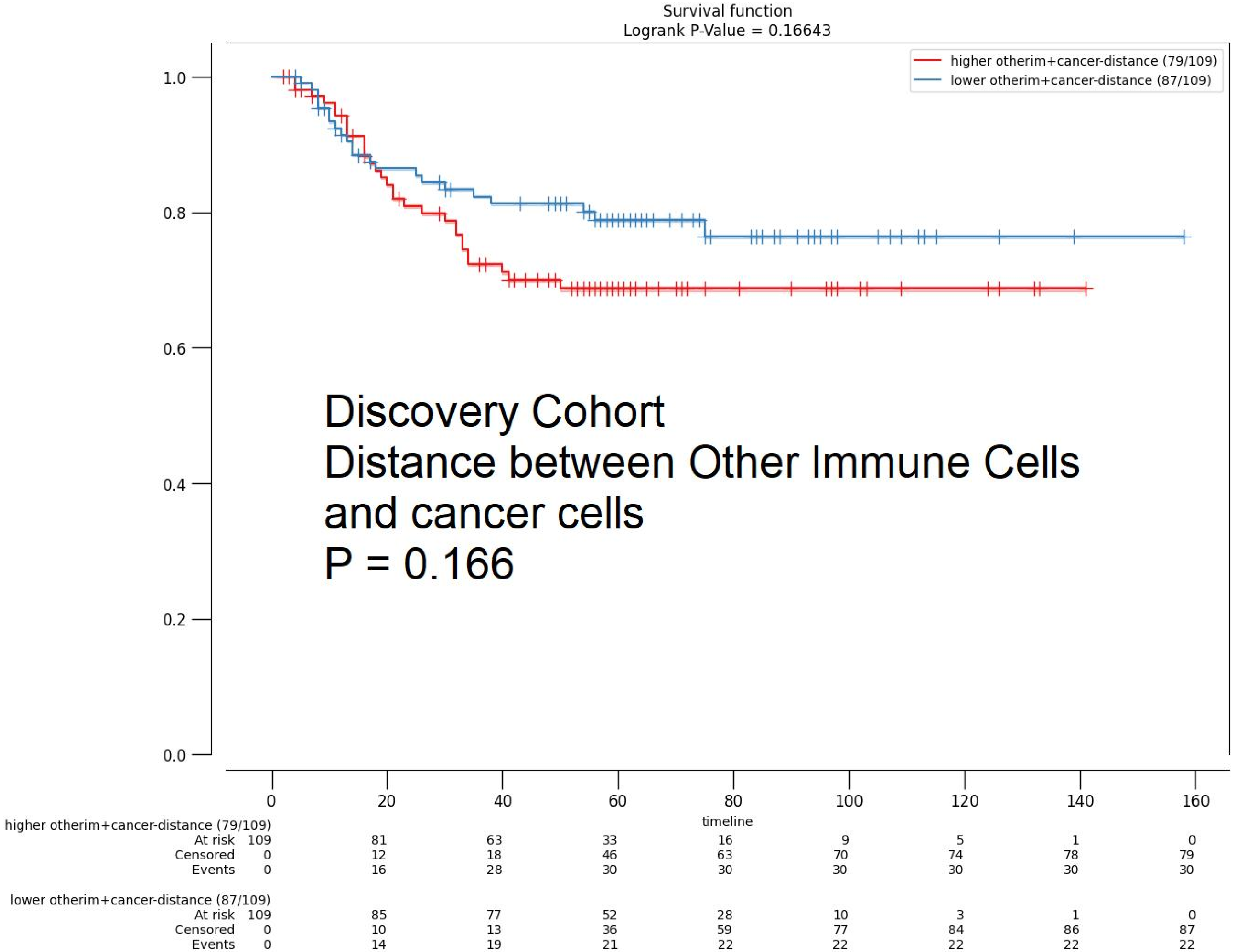
Survival function
Logrank P-Value = 0.16643
higher otherim+cancer-distance (79/109)
lower otherim+cancer-distance (87/109)
1.0
0.8
0.6
0.4
0.2
0.0
Discovery Cohort
Distance between Other Immune Cells
and cancer cells
P = 0.166
0
20
40
60
80
100
120
140
160
timeline
higher otherim+cancer-distance (79/109)
At risk 109 81 63 33 16 9 5 1 0
Censored 0 12 18 46 63 70 74 78 79
Events 0 16 28 30 30 30 30 30 30
lower otherim+cancer-distance (87/109)
At risk 109 85 77 52 28 10 3 1 0
Censored 0 10 13 36 59 77 84 86 87
Events 0 14 19 21 22 22 22 22 22

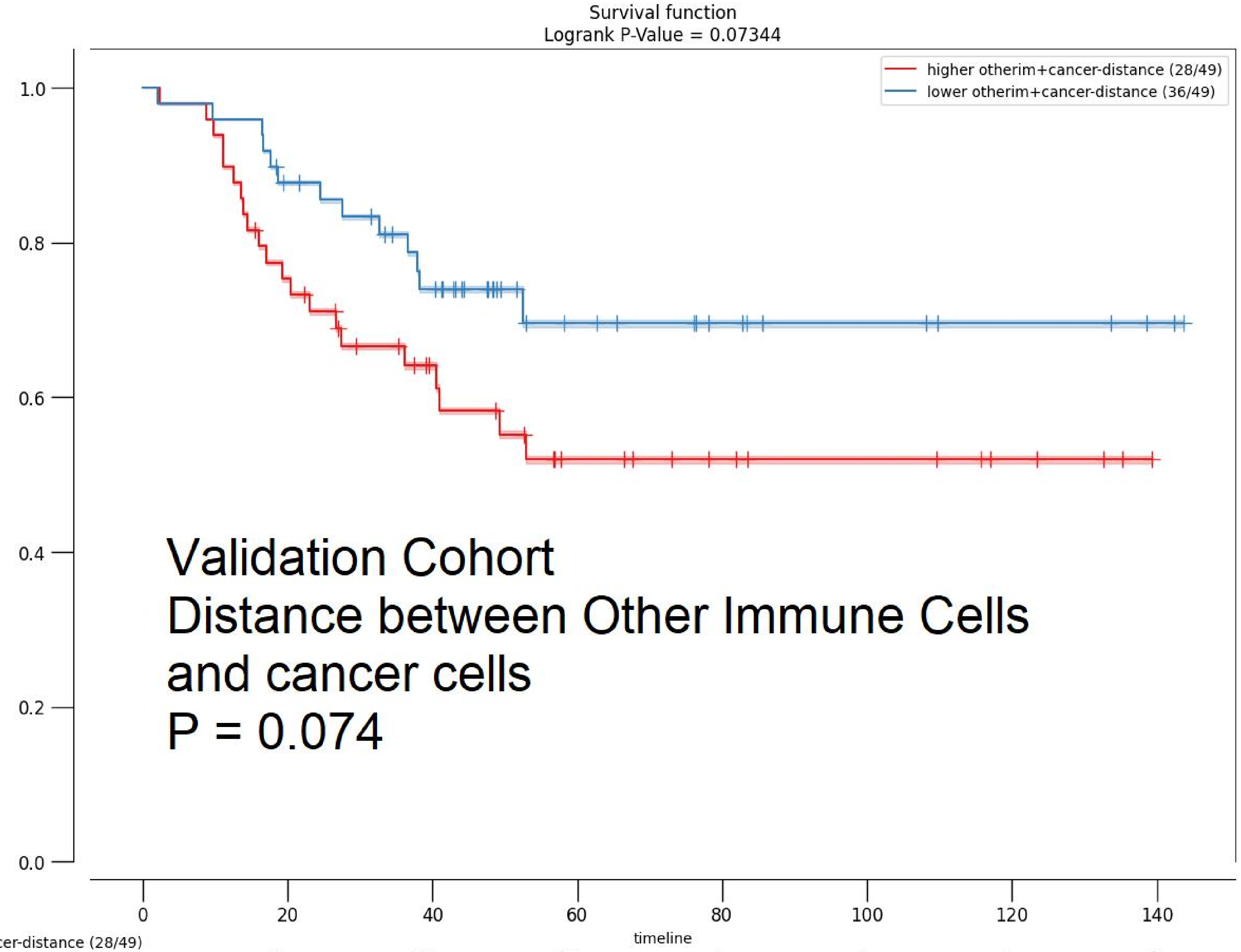


| | 0 | 20 | 40 | 60 | 80 | 100 | 120 | 140 |
|---|---|---|---|---|---|---|---|---|
| **higher otherim+cancer-distance (28/49)** | | | | | | | | |
| At risk | 49 | 36 | 22 | 13 | 9 | 7 | 4 | 0 |
| Censored | 0 | 1 | 10 | 15 | 19 | 21 | 24 | 28 |
| Events | 0 | 12 | 17 | 21 | 21 | 21 | 21 | 21 |
| **lower otherim+cancer-distance (36/49)** | | | | | | | | |
| At risk | 49 | 41 | 31 | 14 | 9 | 6 | 4 | 2 |
| Censored | 0 | 2 | 6 | 22 | 27 | 30 | 32 | 34 |
| Events | 0 | 6 | 12 | 13 | 13 | 13 | 13 | 13 |
